# Global and cell type-specific immunological hallmarks of severe dengue progression

**DOI:** 10.1101/2022.12.11.519930

**Authors:** Luca Ghita, Zhiyuan Yao, Yike Xie, Veronica Duran, Halise Busra Cagirici, Jerome Samir, Ilham Osman, Olga Lucia Agudelo Rojas, Ana Maria Sanz, Malaya Kumar Sahoo, Makeda L. Robinson, Rosa Margarita Gelvez, Nathalia Bueno, Fabio Luciani, Benjamin A. Pinsky, Jose G. Montoya, Maria Isabel Estupiñan Cardenas, Luis Angel Villar Centeno, Elsa Marina Rojas Garrido, Fernando Rosso, Stephen R. Quake, Fabio Zanini, Shirit Einav

**Author notes:** these authors contributed equally (Luca Ghita, Zhiyuan Yao, Yike Xie, Veronica Duran). these authors also contributed equally (Fabio Zanini, Shirit Einav).

## Abstract

Severe dengue (SD) is a major cause of morbidity and mortality impacting approximately 5 million of the 400 million people infected with dengue virus (DENV) annually. To define DENV target cells and immunological hallmarks of SD progression in children’s blood, we integrated virus-inclusive single cell RNA-Seq 2 (viscRNA-Seq 2) with functional assays. Beyond myeloid cells, in natural infection, B cells harbor replicating DENV capable of infecting permissive cells. Alterations in cell type abundance, gene and protein expression and secretion, and cell-cell communications point towards increased migration and inflammation in SD progressors (SDp). Concurrently, antigen presenting cells from SDp demonstrate intact uptake, yet impaired interferon responses and antigen presentation, in part DENV-modulated. Increased activation, regulation, and exhaustion of effector responses and expansion of HLA-DR-expressing, possibly compensatory, adaptive-like NK cells also characterize SDp. These findings reveal DENV target cells in the human blood and provide insight into SD pathogenesis beyond antibody-mediated enhancement.

## Introduction

Dengue virus (DENV) is a mosquito-borne, positive-sense single-strand RNA *flavivirus* estimated to infect 400 million people annually in over 128 countries^1^. Most DENV-infected individuals are asymptomatic or exhibit an acute febrile illness called dengue fever (D). However, 5-20% of symptomatic patients progress within several days of symptom onset to severe dengue (SD), manifesting by hemorrhage, shock, organ failure and sometimes death^2, 3^. For unknown reasons, SD progression in children has been associated with greater plasma leakage and mortality rate than adults^4, 5^.

The identification of SD progressors (SDp) early during the disease course is currently ineffective, relying on broadly defined and nonspecific clinical warning signs that often develop late during the disease course and whose global implementation in 2009 has increased hospitalization rates, challenging resource allocation and not eliminating morbidity and mortality^6, 7^. Moreover, the development of effective dengue vaccines and antivirals to prevent SD progression has been hindered^8^.

A major risk factor for SD is the presence of non-neutralizing anti-DENV antibodies, mediating antibody-dependent enhancement (ADE) upon secondary infection with a heterologous DENV serotype (of four circulating)^9, 10^. Nevertheless, reliance on bulk tissue assays (e.g. whole blood microarrays) and immunocompromised mouse models have somewhat limited characterization of the cellular targets of DENV and the roles of other immune responses in SD pathogenesis in humans. There is therefore an urgent need to better understand SD pathogenesis and identify biomarkers predictive of disease progression and targets to prevent it.

We have previously developed virus-inclusive single-cell RNA-Seq (viscRNA-Seq), an experimental and computational approach to simultaneously detect host and viral transcripts, and used it to investigate molecular signatures of SD progression in 10 peripheral blood mononuclear cell (PBMC) samples from adults^11, 12^. Here, to investigate the cellular and molecular determinants of SD progression in children, we profiled the host and viral transcriptomes in PBMCs derived from 20 DENV-infected children and 4 healthy controls using an optimized viscRNA-Seq 2 approach. We computed the abundance, gene expression and communication network of multiple distinct cell types across disease categories and validated key observations by single-cell proteomics and functional assays in patient-derived samples. Our findings reveal DENV target cells in the human blood and point to key molecular mechanisms preceding SD progression.

## Results

### Host and viral transcripts detected via viscRNA-Seq 2 in subtypes of PBMCs from DENV-infected children

To characterize the immune response associated with SD progression during natural infection in children, we profiled the transcriptome of PBMCs collected early during the disease course. Individuals who presented within 1 to 5 days from symptom onset and tested positive for DENV were enrolled in our Colombia dengue cohort^7, 12^. Disease severity was defined as dengue (D), dengue with warning signs (DWS) and severe dengue (SD)^3^ at presentation and discharge. Here, we studied samples from four healthy donors (H, n=4) and 20 DENV-infected 4-17-year-old children, of whom seven progressed within days following enrollment to SD, one presented with SD (SDp, n=8), and twelve had uncomplicated disease course (DWS, n= 4; D, n=8) (**Fig. 1a, Table 1**).

**Fig.1:**
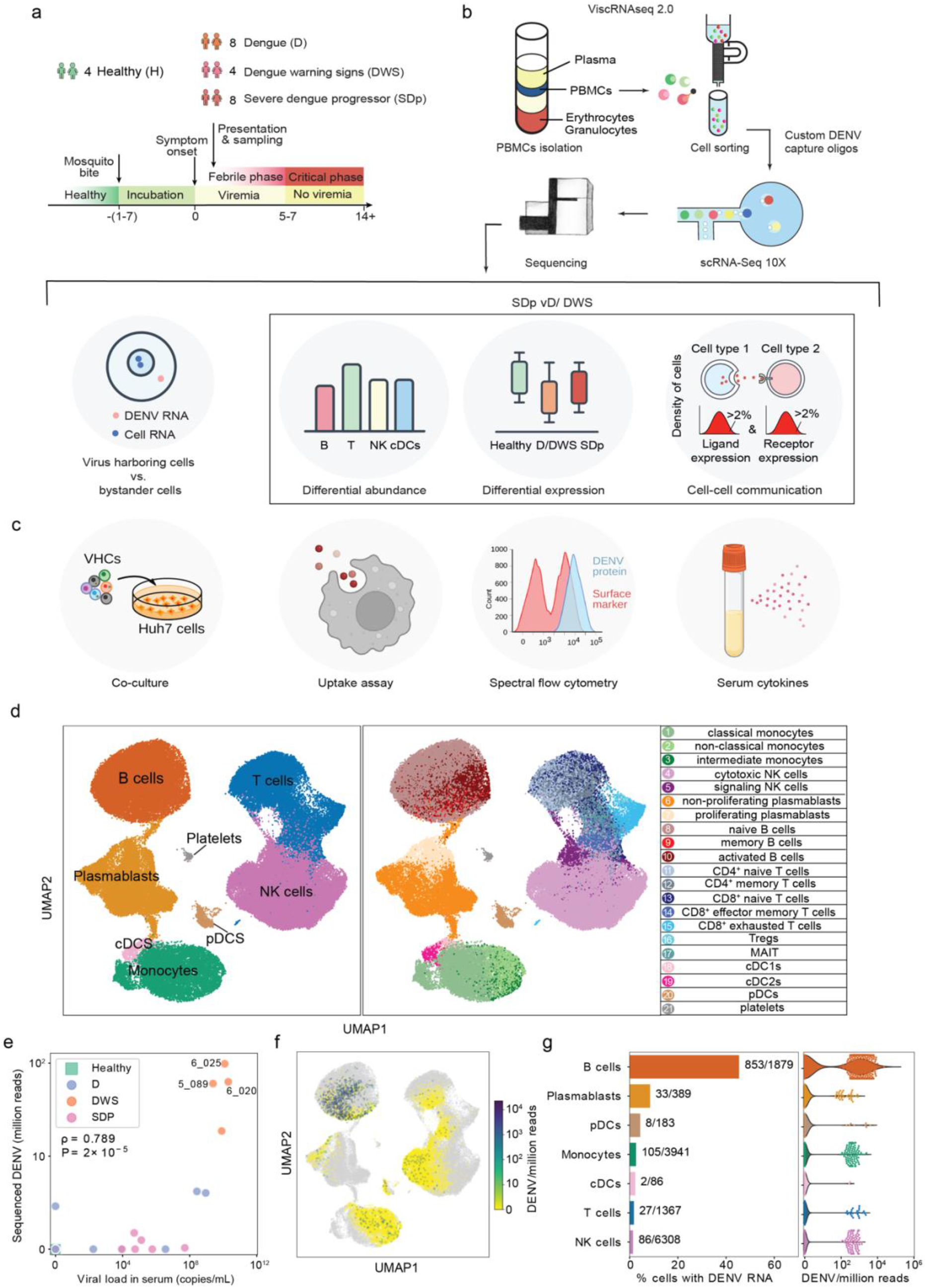
viscRNA-Seq 2 analysis of DENV-infected PBMCs defines 21 immune subtypes and VHCs. **a-c**, Schematic of the study design. **a**, PBMC and serum samples were collected from children enrolled in the Colombia dengue cohort upon presentation to the clinic (D, DWS, SD) and from healthy children (H). **b**, Schematic of the experimental and computational components of viscRNA-Seq 2. **c**, Outline of the performed functional and validation assays on patient-derived PBMCs. **d**, UMAP embedding of the viscRNA-Seq 2 dataset indicating broad cell types (left) and subtypes (right). **e**, Scatter plot of normalized DENV reads measured in PBMCs by viscRNA-Seq 2 versus serum viral load measured by RT-qPCR (Spearman correlation coefficient ρ=0.76, p-value=1.86e-5). Each dot represents a single sample, color-coded by disease severity (H=green; D=blue; DWS=orange; SD progressor=pink). Sample IDs are shown for the 3 DWS patients with highest DENV read counts. **f**, UMAP color-coded by DENV reads per million reads of the 3 patients labeled in **e. g**, Fraction of VHCs from the total (bar plot, left panel) and distribution of DENV read per million reads (violin plots, right panel) in the indicated immune cell types from the 3 patients labeled in **e**.

viscRNA-Seq 2’s improvements we introduced include magnetic cell sorting to more gently enrich for rare cell types, redesigned DENV capture oligos to improve DENV viral RNA (vRNA) capture, and 5’-droplet based microfluidics (10X Genomics) to increase cell numbers (see methods, **Fig. 1b**). This approach enabled simultaneous coverage of transcriptome from 193,727 PBMCs, of which 112,403 were high quality (see methods, **Table 2**). Because of large patient-to-patient sample variability, we annotated cell types and subtypes within each patient using Leiden clustering^13^ (see methods). Overall, 8 cell types and 21 subtypes were identified (**Fig. 1d**) and verified by the expression of unique marker genes (**Extended Data Fig. 1a**).

To define the cell types that harbor vRNA, we used serotype-specific oligonucleotides complementary to the positive strand of the viral genome, detecting over 3,000 vRNA molecules across 1,373 cells from 10 DENV-infected samples across disease categories (**Extended Data Fig. 1b**). vRNA molecule number positively correlated with the corresponding serum viral load (**Fig. 1e**). B cells were the predominant vRNA harboring cells (VHCs), with over 40% of B cells harboring vRNA and showing highest number of DENV reads per cell in three DWS samples (vs. less than 5% of monocytes and NK cells), proposing B cells as DENV targets (**Fig. 1e**,**f**,**g; Extended Data Fig. 1b**,**c**).

### DENV actively replicates in patient-derived B cells and alters their transcriptional profile

Since myeloid cells were reported to be the primary targets of DENV in the blood^14, 15^, yet vRNA was more abundant in B cells, we measured the intracellular expression of DENV envelope (E) protein via spectral flow cytometry in PBMCs (**Extended Data Fig. 1a**) from six viremic DWS patients (**Table 1**). Patient-derived B cells (CD19^+^CD20^+^) expressed moderate, albeit variable, E levels relative to healthy control (**Fig. 2a**,**b**). High E expression levels were measured in classical (CD14^+^CD16^-^), non-classical (CD16^+^) and intermediate (CD14^dim^CD16^+^) monocytes, type II conventional dendritic cells (cDC2) (CD1c^+^), and CD56^bright^ NK cells. E protein expression was low in cDC1s (CD141^+^) and practically absent in pDCs and T cells (**Fig. 2a**,**b**).

**Fig.2:**
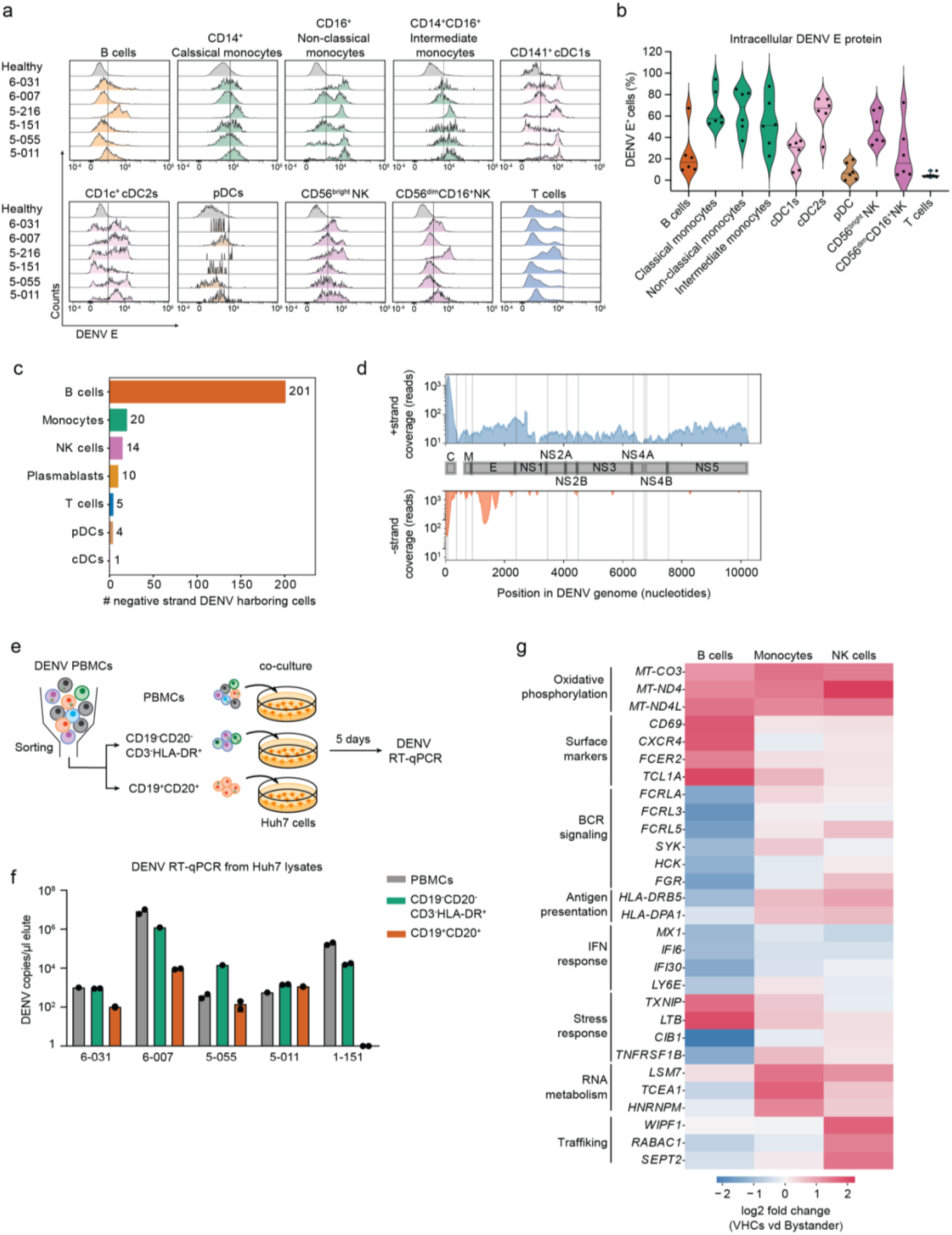
DENV actively replicates in patient-derived B cells and alters their transcriptional profile. **a**,**b** Distributions of intracellular expression of DENV envelope (E) protein (**a**) and % of DENV E positive cells (**b**) in B cells (CD19^+^), classical monocytes (CD14^+^CD16^-^), non-classical monocytes (CD14^-^CD16^+^), intermediate monocytes (CD14^+^CD16^+^), cDC1s (CD141^+^), cDC2s (CD1c^+^), pDCs (CD123^+^CD303^+^), CD56^bright^ NK cells, CD56^dim^CD16^+^ NK cells, and T cells from the indicated DENV-infected patients with viremia ranging from 10^5^ to 10^9^ viral copies/mL (N=2,n=6 combined data) and healthy controls (N=2, n=6 combined data) measured via spectral flow cytometry. **c**, Percentage of negative strand RNA-harboring cells (VHCs) over total number of VHCs across immune cell types in three DWS patients with highest vRNA reads shown in Fig.1e. **d**, Coverage of detected positive (+ strand) and negative (– strand) vRNA strands along the DENV genome. **e**, Schematic of the co-culture experiments shown in f. **f**, Bar plot showing DENV RNA copies per μL measured via RT-qPCR in Huh7 lysates following a 5-day co-culture with patient-derived total PBMCs (gray), or HLA-DR^+^ (CD19^-^CD20^-^ CD3^-^HLA-DR^+^ cells, green) and B cell (CD19^+^CD20^+^, orange) fractions isolated from the same 5 DWS patients (N=2, n=5). **g**, Heatmap showing log2 fold change of DEGs between VHCs and corresponding bystander B cells, monocytes, and NK cells in 3 DWS patients with highest vRNA reads shown in **Fig.1e**. DEGs were identified by the median log2 fold change of 100 comparisons between VHCs and equal numbers of random corresponding bystander cells. N, number of experiments conducted; n, number of samples per experiment.

To determine whether DENV actively replicates in B cells during natural infection, we leveraged the ability of viscRNA-Seq 2 to ascertain for each vRNA molecule which genomic strand it originated from. We detected 419 DENV reads across 255 cells that originated from the DENV negative strand (presumably synthesized during vRNA replication) (**Fig. 2c**).

These molecules were detected primarily in B cells and distributed at the 5’ end of the genome, similarly to positive-strand reads, and at position ∼1700 bases—a region that does not contain tight RNA hairpins^16^ possibly providing easier template access for the poly-T oligonucleotide (**Fig. 2c**,**d**). The ratio of negative-strand to total vRNA was 14.1%, an order of magnitude higher than the ratio of antisense molecules for representative housekeeping genes (**Extended Data Fig. 2b**), favoring active DENV replication over an artifact of library preparation.

To assess the ability of patient-derived B cells to produce infectious virus particles and infect naive cells, we co-cultured DENV-permissive human hepatoma (Huh7) cells for 5 days with PBMCs from 5 viremic patients or with two cell fractions isolated from the same samples via flow cytometry: i) B (CD19^+^CD20^+^) cells; and ii) HLA-DR^+^ cells (CD45^+^CD3^-^CD19^-^CD20^-^HLA-DR^+^), followed by RT-qPCR measurement of intracellular DENV copies number (**Fig. 2e, Extended Data Fig. 2c, Table 1**). High DENV copy numbers were measured in co-cultures of PBMCs from all 5 patients, indicating effective infectious virus production (**Fig. 2f**). Isolated HLA-DR^+^ cells from 5 out of 5 patients effectively infected Huh7 cells with DENV copy numbers that were comparable or slightly lower than PBMCs. Remarkably, sorted B cells from 4 out of the 5 patients infected Huh7 cells, with DENV copy numbers that were comparable or up to 2-fold lower than the PBMC and HLA-DR^+^ fractions (**Fig. 2f**). DENV copy numbers in the various co-cultures positively correlated with viremia level (**Extended Data Fig. 2d**).

To determine whether DENV alters the transcriptome of VHCs, we identified differentially expressed genes (DEGs) between vRNA-harboring and the corresponding bystander B cells, monocytes, and NK cells using a bootstrapping strategy (see Methods) (**Fig. 2g**). Oxidative phosphorylation genes were upregulated in VHCs relative to bystanders in all three cell types, suggesting DENV-induced intracellular stress. vRNA-harboring B cells showed upregulation of genes encoding surface markers (CD69, CXCR4) and downregulation of genes involved in BCR signaling (*FCRLA, FCRL3, FCLR5, HCK, SYK, FGR*), antigen presentation (*HLA-DPA1, HLA-DRB5)*, and interferon stimulated genes (ISGs) (*MX1, IFI6, IFI30*, L*Y6E*).

These results reveal infectious DENV production in natural infection in circulating B cells, in addition to monocytes and cDCs^14, 15^, associated with a naive cell state and altered antigen presentation and interferon signaling signatures in dynamic balance with bloodstream virions.

### Alterations in cell subtype abundance detected via viscRNA-seq 2 distinguish SDp from uncomplicated dengue patients

To assess whether SD progression is associated with alterations in cell abundance, we compared the composition of cell subtypes within the major immune cell populations in SDp, symptomatic non-progressors (D, DWS), and healthy controls (**Fig. 3a**). The fractions of classical monocytes, signaling NK cells and proliferating plasmablasts were significantly expanded in SDp relative to non-progressors. A Treg population was present only in SDp, and the abundance of CD4^+^ memory and CD8+ exhausted T cell fractions was higher in SDp than non-progressors, albeit these differences were not statistically significant (**Fig.3a**,**b**). In contrast, smaller fractions of non-classical monocytes, cytotoxic NK cells and non-proliferating plasmablasts were found in SDp relative to non-progressors (**Fig. 3a**). These differences in cell subtype abundance did not correlate with serum viral load (**Extended Data Fig. 1a**).

**Fig.3:**
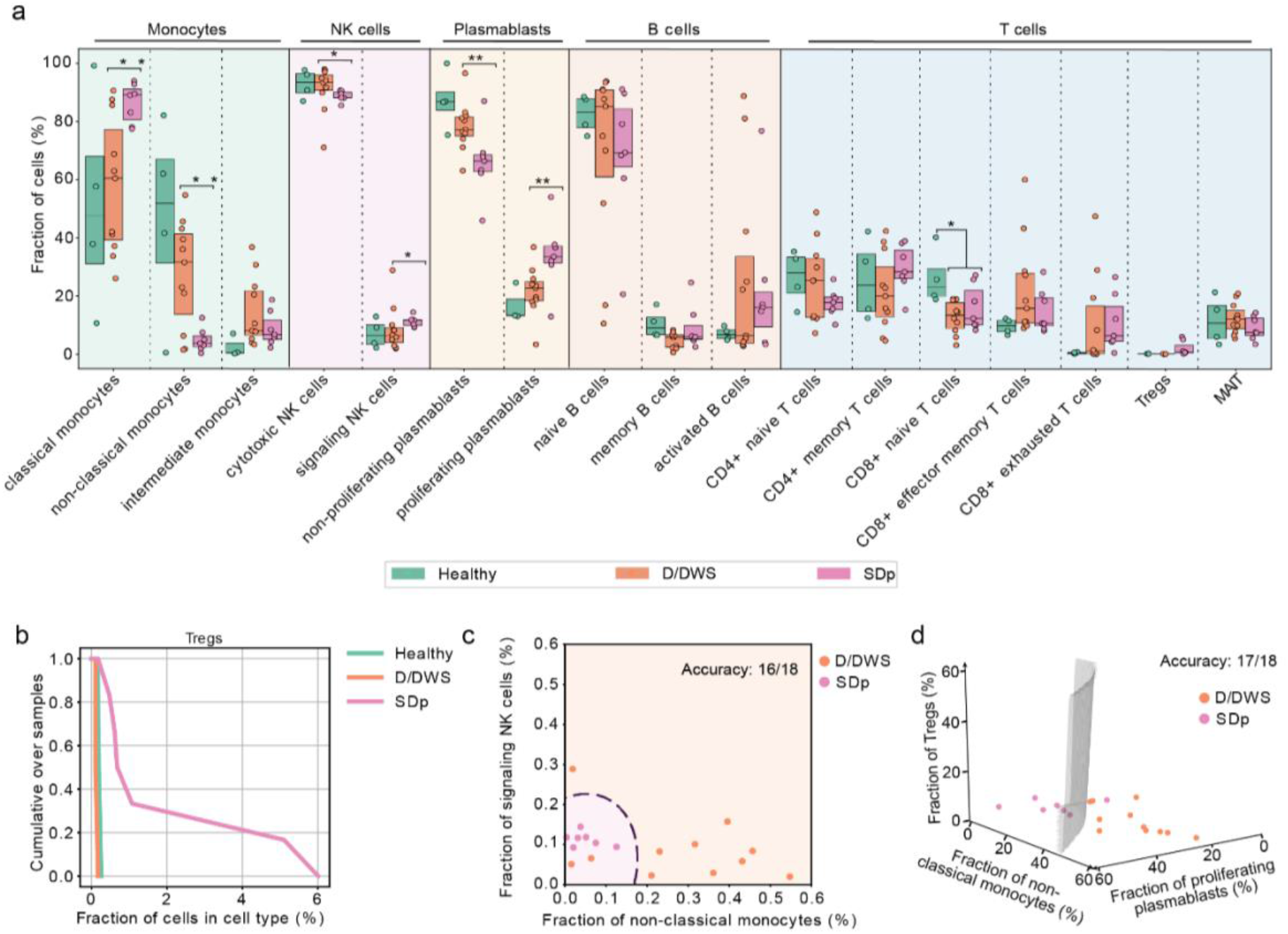
Cell subtype abundance detected via viscRNA-Seq 2 classifies DENV disease categories. **a**, Box plots showing the fractions (%) of cell subtypes within each major immune cell type (*p-value<0.05, **p-value<0.001 by ks test). Each dot represents an individual, color-coded by disease severity (H=green; D=blue; DWS=orange; SDp=pink). Box plots’ horizontal lines indicate the first, second (median) and third quartiles. **b**, Cumulative distribution of Treg fractions within T cells over all patients across disease severity. **c**,**d** Two (c)- and three (d) dimensional Support Vector Machine (SVM) classifiers for SDp versus D/DWS using the fraction of cells indicated on the axes. Accuracy is evaluated using leave-one-out cross-validation.

To determine whether changes in cell subtype abundance could classify SDp from D and DWS patients, we applied a supervised machine learning algorithm (see methods). Using solely the abundance of signaling NK cell and non-classical monocyte fractions as discriminatory parameters effectively distinguished SDp in most leave-one-out comparisons (16 out of 18, **Fig. 3c**), and addition of Treg abundance improved the classification (17 out of 18 correct, **Fig. 3d**).

These findings propose that altered immune regulation characterized by increased pro-inflammatory cells (increased CD16 monocytes and proliferating plasmablasts) yet suppressed effector cells (decreased cytotoxic NK cells, Treg emergence) is associated with SD progression.

### Antigen presenting cells (APCs) from SDp show increased FCγ-receptor and uptake activity yet decreased antigen presentation signatures

We profiled the transcriptional landscape of APCs associated with SD progression via pairwise differential expression analysis between individual SDp and D patients (see methods, **Extended Data Fig. 4a, Table 3**). Gene ontology (GO) analysis of differentially expressed genes (DEGs) in monocytes revealed upregulation of pathways associated with pro-inflammatory and myeloid cell activation responses and downregulation of pathways associated with cytokine-mediated signaling, type I and type II interferon response, and antigen processing and presentation in SDp relative to D (**Extended Data Fig. 4b**).

DEGs between SDp and D were also identified in the three monocyte subtypes and in other APCs: cDCs and B cells. Among the upregulated genes in monocytes were pro-inflammatory genes, FCγ-receptor signaling genes (*FCGR1A, FCGR2A, PRKCD, BTK, ITPR2*), the scavenger receptor *CD163*, and genes involved in cell adhesion and migration, with greater upregulation of most genes in intermediate than classical and non-classical monocytes (**Fig. 4a**). Pro-inflammatory and cell adhesion and migration genes were also upregulated in cDCs from SDp relative to D, with variable expression patterns in cDC1s and cDC2s (**Fig. 4b**). B cells from SDp showed upregulation of genes related to host and viral translation, RNA binding and differentiation (**Fig. 4c**).

**Fig.4:**
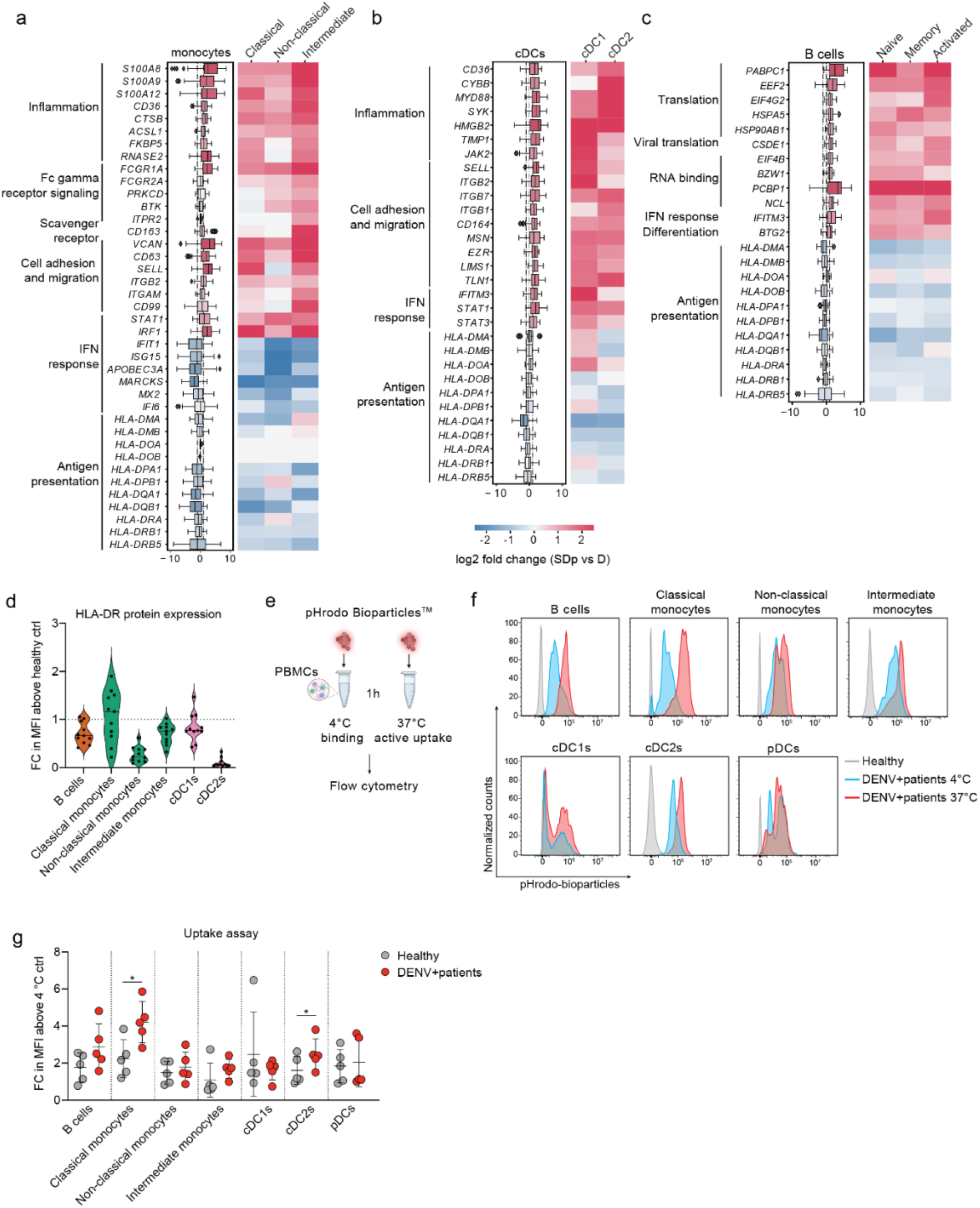
Immunological determinants of SD progression in patient-derived antigen presenting cells (APCs). **a-c**, Differentially expressed genes (DEGs) between D and SDp detected in monocyte (a), cDC (b) and B cell (c) populations (Box plots, left) and the corresponding distinct cell subtypes (heatmaps, right). Data are color-coded based on median log2 fold change of pairwise comparisons. Box plots’ horizontal lines indicate the first, second (median) and third quartiles. **d**, Violin plot showing HLA-DR cell surface expression measured via spectral flow cytometry in patient-derived PBMCs expressed as fold change (FC) of mean fluorescence intensity (MFI) above healthy control (n=11, N>2). Dotted line represents mean value of healthy controls normalized at value 1, horizontal lines in violin plots represent median. **e**, Schematic of the uptake experiments shown in f and g. Patient-derived PBMCs were incubated for 1 hour either at 4°C or 37°C with pHrodo Bioparticles followed by fluorescence measurement via spectral flow cytometry. **f**, Histograms showing the percentage of pHrodo Bioparticles positive cells in distinct cell subtypes derived from healthy (gray) and DENV-infected patients measured at 4 °C (cyan) or 37 °C (red). **g**, Bioparticle uptake quantification in the indicated cell subtypes shown as FC of mean MFI above 4 °C control. Dots represent individual patients, color-coded by disease status: healthy (gray) and DENV-infected (red) (n=5, N>2) (* = p<0.05, two-tailed Mann-Whitney test).

Interestingly, whereas interferon signaling genes, such as *IRF1* and *STAT1*, were upregulated in all APC subtypes in SDp vs. D, ISGs, such as *IFIT1, ISG15, MARCKS* and *APOBEC3A* where upregulated in cDCs and B cells but downregulated in monocytes, particularly non-classical monocytes, suggesting that progression to SD may be associated with dysregulated IFN response (**Fig. 4a-c**).

Multiple MHC class II genes were universally downregulated in all APC subtypes in SDp vs. D (**Fig. 4a-c**). Lower expression of HLA-DR protein was also detected by flow cytometry analysis on DENV-infected patient-derived APCs (except for CD14^+^ monocytes) than healthy (**Fig. 4d**), confirming our recent mass cytometry (CyTOF) findings^17^ and proposing decreased antigen presentation capacity as another phenotype of SD progression.

Compellingly, this finding was associated with upregulation of F*CGR1A* (CD64), *FCGR2A* (CD32) and *CD163* in monocytes from SDp vs. D (**Fig. 4a**) and in the combined DENV-infected population vs. healthy (**Extended Data Fig. 4c**). Flow cytometry analysis confirmed the latter at the protein level, showing increased expression levels of CD64 (in agreement with our recent CyTOF data^17^) and CD163 (albeit variable expression of CD32) on APC subsets in 6 additional DENV-infected patients vs. healthy (**Extended Data Fig. 4d**). To functionally profile the phagocytic ability of APCs, we measured the uptake activity of these patient-derived APCs using a pHrodo bioparticles assay either at 4 °C (binding control) or at 37 °C (active uptake) followed by flow cytometry analysis. A 2-4-fold increase in mean fluorescence intensity (MFI) above the binding control, indicative of increased uptake, was measured in 5 DENV-infected relative to 5 healthy individuals, both globally (**Fig. 4f**) and patient by patient (**Fig. 4g**), in classical (CD14^+^CD16^-^) monocytes, cDC2s (CD1c^+^), and B cells (CD19^+^CD20^+^), albeit the latter was not statistically significant.

These findings provide evidence that SD progression is associated with an inflammatory phenotype accompanied by impaired interferon response and antigen presentation in APCs despite intact uptake activity.

### SD progression is associated with increased activation and exhaustion of effector cells

We extended the differential expression analysis between SDp and D to effector cells (**Table 3**). In NK cells, GO analysis of DEGs revealed upregulation of T cell proliferation or activation and IFNγ response and downregulation of chemotaxis, migration and cytokine signaling pathways (**Extended Data Fig. 1a**). Both cytotoxic and signaling NK cells demonstrated upregulation of genes involved in cytoskeleton rearrangement and protein translation and folding in SDp (**Fig. 5a**). Cytotoxic NK cells upregulated MHC-II genes (*HLA-DRB1, HLA-DPA, HLA-DPB1*), pro-inflammatory (*CCL5* and *IL32)*, and cell migration genes ..CD2 (cytotoxicity activator) and downregulated *KLRB1* (cytotoxicity suppressor) (**Fig. 5a**).

**Fig.5:**
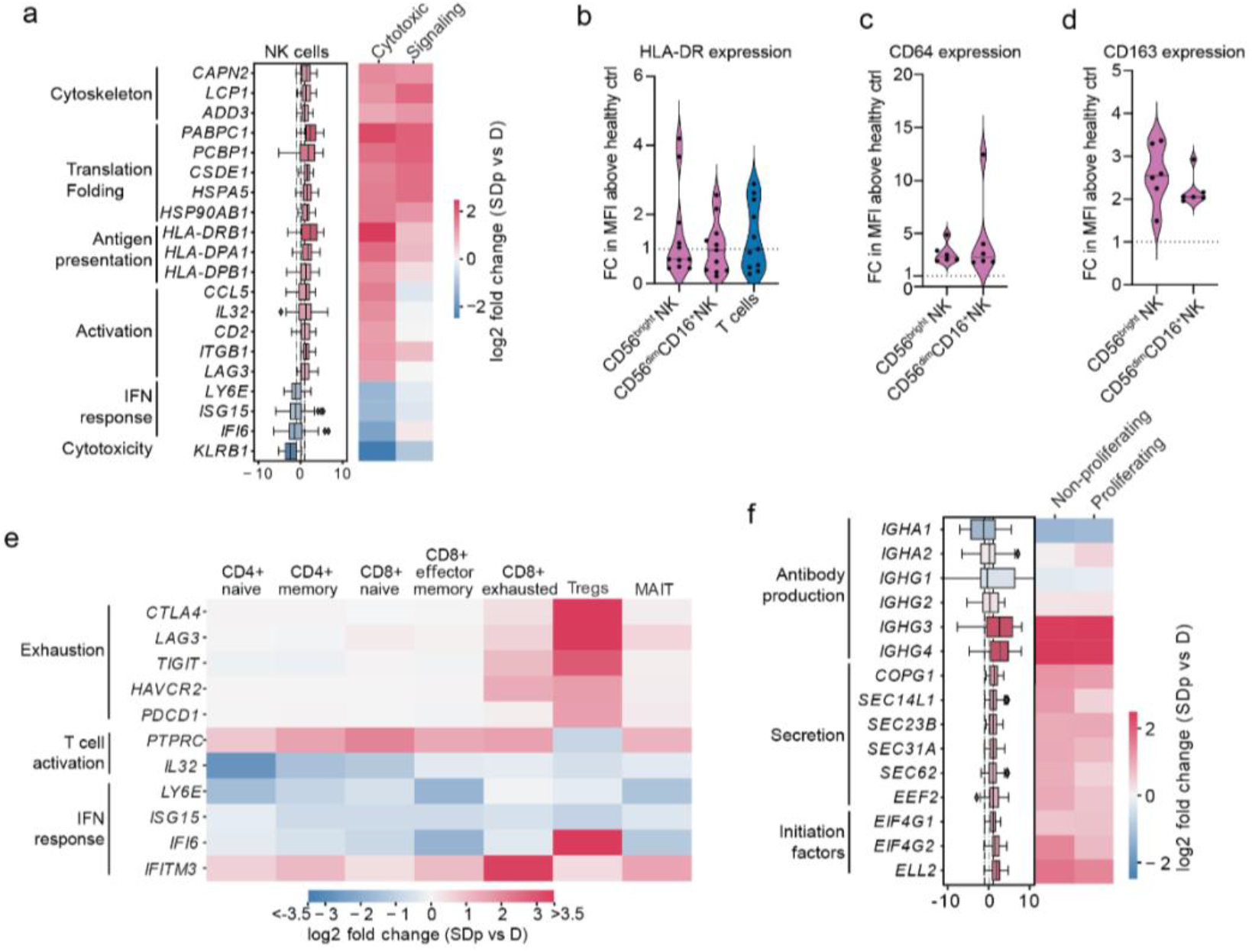
Phenotypic alterations in effector cells associated with SD progression. **a, e, f**, DEGs between D and SDp detected in NK cell (a) and plasmablast (e) populations (Box plots, left) and distinct NK cell, T cell (f) and plasmablast subtypes (heatmaps, right). Data are color-coded based on median log2 fold change of pairwise comparisons. Box plots’ horizontal lines indicate the first, second (median) and third quartiles. **b-d**, Violin plots showing cell surface expression of HLA-DR (b) (n=11, N>2), CD64 (c) (n=6, N=2) and CD163 (d) (n=6, N=2) measured in patient-derived NK (b-d) and T cells (b) via spectral flow cytometry expressed as fold change (FC) of mean fluorescence intensity (MFI) above healthy control. Dotted line represents mean value of healthy controls normalized at value 1, horizontal lines in violin plots represent median.

This signature, suggestive of cytotoxic NK cell activation, was accompanied by upregulation of *LAG3*, an exhaustion marker, and downregulation of ISGs (*LY6E, ISG15, IFI6)* (**Fig. 5a**).

In agreement with the transcript data and in contrast with classical APCs (**Fig. 4**), the expression of HLA-DR protein measured by flow cytometry was higher in DENV-infected, patient-derived CD56^bright^ and CD56^dim^CD16^+^ NK cells than healthy controls (**Fig. 5b**). Moreover, the abundance of *HLA-DRA* positive NK cells was greater in SDp than D (**Extended Data Fig. 5b**). These findings were associated with higher expression of CD64, CD32, and CD163 proteins measured by flow cytometry in patient-derived NK (particularly cytotoxic) than healthy, yet no alterations in uptake activity were observed (**Fig.5b**,**c; Extended Data Fig. 5c**,**d**).

Evidence for exhaustion and regulation in SDp was identified also in T cells, particularly exhausted CD8^+^ T cells and Tregs, cell subtypes whose abundance increased in SDp (**Fig. 3a**). These cells demonstrated upregulation of *CTLA4, LAG3, TIGIT, HAVCR2*, and *PDCD1* (**Fig. 5d**), in agreement with increased CTLA4 and PD-1 measured on Tregs via CyTOF^17^. Notably, these findings were associated with signatures suggestive of attenuated effector functions across most T cell subtypes (**Fig. 5d**). For example, *IL32*, whose expression typically increases upon T cell activation to induce pro-inflammatory cytokine production in myeloid cells, was downregulated in SDp. Moreover, multiple ISGs, including *LY6E, ISG15*, and *IFI6*, but not *IFITM3*, were downregulated in SDp vs. D.

The transcriptome of plasmablasts was characterized by upregulation of specific immunoglobulin constant region genes (*IGHG3 and IGHG4*) and genes involved in protein translation and secretion in SDp vs. D, and was comparable between proliferating and non-proliferating plasmablasts (**Fig. 5e**).

These findings provide evidence that SD progression is associated with activation and regulation of effector lymphocyte responses and the presence of cytotoxic NK cells that may be taking on an antigen-presenting role.

### SD progression is associated with altered cell-cell communications and cytokine production

Since the immune response is orchestrated by physical and paracrine cell-cell communications facilitated by interactions of cell-surface receptors with membrane-bound or secreted ligands, respectively, we computationally analyzed the cell-cell communications in SDp and D children. We identified 1164 genes encoding proteins involved in known interactions (interacting genes)^18, 19^. A larger number of candidate interactions, most commonly involving monocytes and cDCs, was detected in SDp than D (see methods) (**Fig. 6a**; **Extended Data Fig. 1a**,**b**).

**Fig.6:**
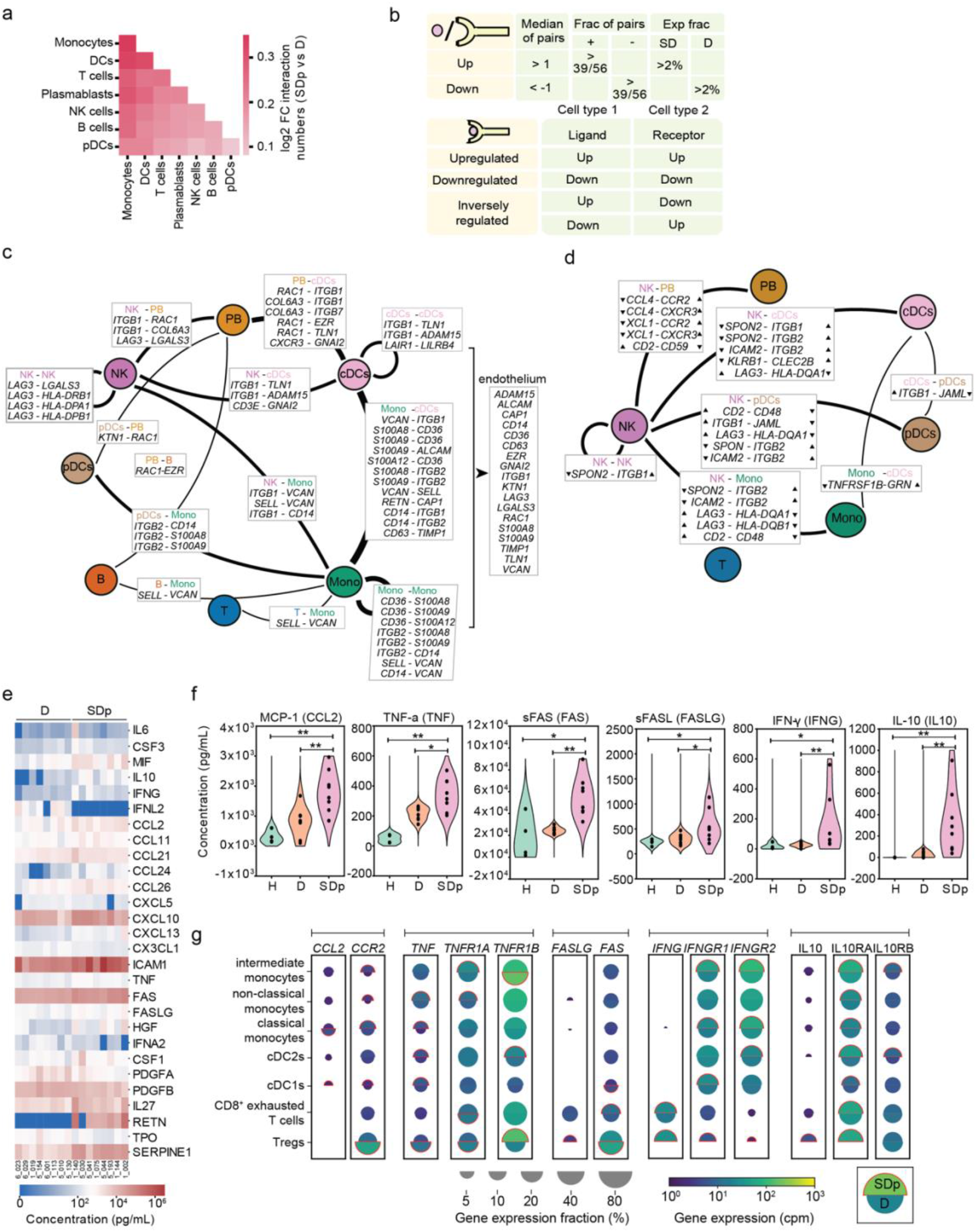
Cell-cell communication and cytokine production are increased in SDp. **a**, Heat map showing the log2 fold change in number of candidate interactions between SDp and D children defined by expression of both the ligand and receptor in at least 2% of cells within each cell type. **b**, Schematic showing the parameters for identifying interacting partners and (top), and the classification of DEIs (bottom). **c, d**, Interaction networks of upregulated (c) and inversely regulated (d) DEIs in SDp vs. D. Circles indicate cell types; lines indicate interaction partners with thickness representing the number of interactions; text boxes specify genes involved in identified candidate interactions and the cell types expressing them, including candidate interactions with endothelium based on Tabula Sapiens data (right panel in c); and arrowheads (in d) depict up- and down-regulation of the indicated genes. **e**, Heatmap showing serum concentrations (pg/mL) of 27 differentially secreted cytokines (out of 80 tested) between SDp vs. D (p-value<0.05, two-tailed Mann-Whitney test). **f**, Violin plots of serum concentration (pg/mL) of top differentially secreted cytokines in SDp vs. D (*p-value<0.05, **p-value<0.001, two-tailed Mann-Whitney test). **g**, Split dot plots showing the expression level of transcripts of cytokines (corresponding to those in (f)), and cognate receptors measured via viscRNA-Seq 2. Top half of each spilt dot plot depicts the expression level in SDp; bottom half depicts the expression level in D; size depicts the percentage of gene expressing cells in the cell types indicated on the left; and color depicts gene expression level in counts per million (cpm). Red outline indicates the condition with the higher gene expression.

We defined interacting DEGs (iDEGs) as ligand- or receptor-encoding genes that were also differentially expressed, and differentially expressed interactions (DEIs) as interactions whose ligand and receptor were iDEGs and passed a statistical randomization test (**Fig. 6b; Extended Data Fig. 6c-d**). DEIs were classified as upregulated, downregulated or inversely regulated according to the regulation of the ligand and receptor. No strongly downregulated DEIs were identified, and the majority of upregulated and inversely regulated DEIs were physical (**Table 4**). iDEGs were abundant and largely upregulated in cDCs and monocytes, and abundant, yet variably regulated in NK cells (**Extended Data Fig. 6e**). Concurrently, cDCs and monocytes formed the largest hubs of the upregulated DEI network (**Fig. 6c; Extended Data Fig. 6c**). Candidate upregulated DEIs were predominantly involved in adhesion and migration (e.g. *ITGB1-VCAN; SELL-VCAN*) and pro-inflammation (e.g. *S100s-CD36*; *RETN-CAP*) (**Fig. 6c; Extended Data Fig.6f**). Eighteen of 31 upregulated iDEGs involved in DEIs in SDp were also expressed in >5% of endothelial cells from the human cell atlas^19^, suggesting their potential role in immune-endothelium interaction (**Fig. 6c**).

In agreement with their involvement in variably regulated iDEGs, NK cells were the hub of the network of inversely regulated iDEGs (**Fig. 6d**,**e; Extended Data Fig. 6d**). Among the candidate communications were: downregulated *SPON2* in NK cells with upregulated *ITGB1* and *ITGB2* in myeloid subtypes; upregulated *LAG* in NK cells with downregulated *HLA-DQA1/HLA-DQB1* in myeloid subtypes, and downregulated *CCL4* and *XCL1* in NK cells (**Fig. 6d; Extended Data Fig. 6f-h**), supporting inhibition of NK cell function and tissue migration.

To strengthen our understanding of paracrine cell-cell communication, we measured the levels of 80 secreted factors in the sera of the same patients and revealed 27 differentially secreted cytokines between SDp and D (**Fig. 6e, Table 5**). Computational analyses revealed specific myeloid and T cell subtypes that may contribute to their secretion and/or interact with these cytokines (**Fig. 6f**,**g**). Increased CCL2 secretion in SDp was associated with upregulation of transcripts for both *CCL2* in cDC1s and its receptor, *CCR2*, in myeloid subtypes, yet downregulation of *CCR2* in Tregs. Moreover, increased TNF, FAS, and FASLG secretion in the sera of SDp suggested apoptosis induction, yet the viscRNA-Seq data revealed distinct cell-type specific responses, with monocytes and cDCs demonstrating proapoptotic signatures (upregulation of *TNF* and *TNFRSF1A*,) vs. Tregs demonstrating antiapoptotic signatures (downregulation of *TNF, TNFRSF1A* and upregulation of the anti-apoptotic *TNFRSF1B*). Increased IFN γ secretion was associated with *IFNG* upregulation in Tregs and exhausted T cells accompanied by upregulation of *IFNGR1* and *IFNGR2* on multiple APCs. Furthermore, an increased secretion of IL10 appears to be partially driven by Tregs and was associated by upregulation of *IL10RA/B* on myeloid cell subtypes and regulatory and exhausted T cells.

SD progression is thus linked to increased coordinated interactions involving myeloid cells associated with migration and pro-inflammation, yet aberrant orchestration of the innate-adaptive immune junction by uncoordinated NK cell interactions.

### A systems immunology model of SD progression

Integrating viscRNA-Seq 2 with functional assay data, we propose a systems immunology model of SD progression in children centered around a disjointed immune response (**Fig. 7**). Whereas uncomplicated D/DWS is characterized by a multifaceted immune response that balances antigen recognition, uptake, and presentation with initiation of DENV-specific responses driven by T and B lymphocytes, SD progression is linked to increased plasmablasts and coordinated interactions promoting migration and pro-inflammation in myeloid cells, yet aberrant orchestration of the innate-adaptive immune junction due to suppressive Tregs and uncoordinated NK cell interactions, promoting tissue injury and reducing viral clearance.

**Fig. 7:**
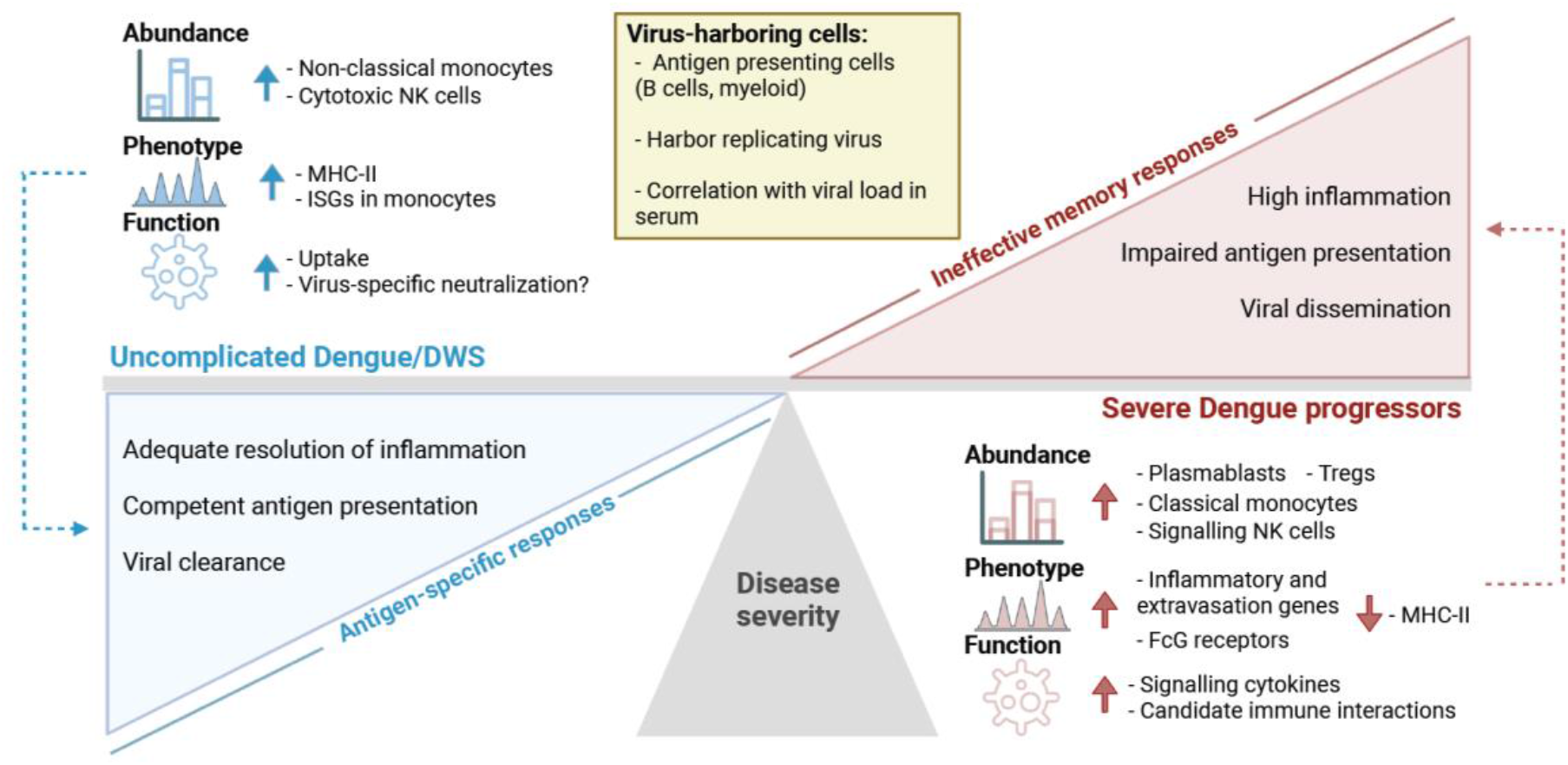
Immune hallmarks of early SD progression. Schematic representation depicting the differences in cell type abundance, phenotype, and immunological function of distinct cell subtypes in SDp (red) vs. uncomplicated D/DWS (blue), and specific attributes of virus-harboring cells (VHCs) (yellow). We propose that B cells and myeloid antigen-presenting cells harbor replication-competent DENV in human blood, and that ineffective memory responses, accompanied by defective antigen presentation and inflammation resolution mechanisms, prevail in SDp, whereas adequate presentation of antigens and effective antigen-specific responses characterize D/DWS patients.

## Discussion

The pathogenesis of DENV infection in humans remains poorly understood, hindering the discovery of biomarkers to predict disease progression and targets to prevent it. Here, we developed viscRNA-Seq 2, a novel scalable approach to capture host and viral reads from ex vivo samples, and combined it with spectral flow cytometry, cytokine and vRNA measurements in serum and functional experiments to discover the cellular targets of DENV in the human blood and immunological determinants of SD progression in children.

Previously, myeloid cells have been reported as the target cells of DENV in natural infection: DENV proteins were detected in cDCs^20^ and monocytes^21^ in patients’ blood via flow cytometry and in macrophages, monocytes and cDCs in clotted blood and tissues from dengue cadavers by immunohistochemistry^22, 23^. These cells are also susceptible to DENV infection in vitro^3, 20^. Here, we not only show that B cells harbor the majority of positive-strand vRNA in children’s blood, as we and others have reported in adult patients^12, 24^, but also measure active replication in myeloid and B cells via three orthogonal experimental methods: negative-strand vRNA detection in patient cells, E protein expression measurement via flow cytometry, and viral spread detection upon co-culture of with permissive cells. Evidence for infectious DENV production was previously demonstrated only in healthy donor-derived B cells in vitro infected at a high multiplicity of infection (MOI=20)^25^, but not in patient-derived B cells. Increased production of subgenomic flavivirus RNAs (sfRNAs) might account for the greater vRNA abundance in B cells yet greater E protein expression in myeloid cells since sfRNAs can (i) suppress antiviral IFN response^26, 27^ consistent with our finding that vRNA-harboring B cells downregulate IFN response genes relative to bystander cells; and (ii) compete for packaging with DENV genomic RNA^28^, increasing intracellular vRNA. Interestingly, vRNA-harboring B cells upregulate transcripts for CD69 and CXCR4—markers for antigen internalization and homing to germinal centers for antigen presentation, respectively—and naive markers. It is intriguing to speculate that by internalizing vRNA, these B cells are triggered towards differentiation without recognizing cognate antigens via their specific BCR and prematurely home to secondary lymphoid organs, thereby expanding plasma cells expressing low-affinity antibodies, the hallmark of ADE. It remains to be determined whether the detection of viral elements in NK cells represents active replication or uptake of infected cells.

While patient-derived monocyte subtypes demonstrate comparable expression of DENV E protein levels and upregulation of genes and ligand-receptor interactions promoting inflammation and migration in SDp, their abundances are altered differently. Classical monocytes are expanded and activated, possibly contributing to SD pathogenesis by promoting migration to lymphoid organs, conferring sustained inflammation and forming a niche for DENV replication, as suggested by others^29^. Contrastingly, the fraction of non-classical monocytes—cells implicated in patrolling the endothelium and protecting from protease-mediated damage^29^—is drastically reduced, potentially impairing immune surveillance and contributing to the endothelial damage observed in SD^30^.

APCs downregulate MHC-II genes involved in antigen presentation in SDp vs. D and reduce cell surface expression of HLA-DR protein in DENV-infected patients vs. healthy, as also observed via CyTOF^17^. Combined with downregulation of ISGs observed in monocytes from SDp relative to D patients, these findings suggest that APCs might fail to adequately trigger antigen-specific T cell-responses, pointing to a lack of coordination between innate and adaptive immunity during early SD progression. Conversely, antigen uptake is greater in all APC subsets, particularly classical monocytes and cDC2s, from DENV-infected patients than healthy donors, possibly compensating for the deficient antigen presentation.

A population of Tregs upregulating transcripts for CTLA4, LAG3, TIGIT, HAVCR2 and PDCD was detected in SDp, suggesting a suppressive phenotype in contrast with another study suggesting reduced suppression^31^, yet in agreement with the detection of CTLA4 and PD-1 overexpressing Tregs via CyTOF analysis of longitudinal samples from our Colombia cohort^17^. Notably, the abundance of Tregs declined prior to SD onset, supporting protection^17^ possibly mediated in part by inhibition of vasoactive cytokine production^32^. However, Treg activation may suppress immune responses, such as via IL-10 secretion^33^, and may thus play a dual role in SD pathogenesis.

Our findings point to NK cells as a key correlate of SD progression. Specifically, cytotoxic NK cells exhibit signatures suggestive of activation based on increased transcripts for CD2 and protein expression of CD64 and CD32, and lower expression of the inhibitory receptor KLRB1 and of CD16 which is cleaved upon NK cell activation^34^. Yet, their abundance is markedly decreased in the circulation, and they demonstrate increased exhaustion based on increased LAG3 expression in SDp, and PD-L1 expression detected by CyTOF^17^, suggesting reduced killing of DENV-infected cells. Alternatively, since CD16 and IFNγ- mediated CD64 expression on NK cells have been implicated in antibody-dependent cellular cytotoxicity (ADCC), one mechanism of infected-cell killing^35, 36, 37^, this expression pattern combined with the higher IFNγ secretion we detected in serum of SDp than D, suggest that ADCC may be induced in SDp.

A subset of cytotoxic NK cells whose abundance is increased in SDp may take an antigen-presenting role based on increased expression of MHC class II transcripts and proteins, IL32, CD2, and the exhaustion marker LAG342 expression, and intact uptake activity. It is tempting to speculate that these adaptive-like NK cells may activate antigen-specific T cell responses, as shown in other disease models^38^, to compensate for the detected impairment in SDp. Concurrently, we identified expansion of signaling NK cells with a pro-inflammatory and migratory phenotype, possibly contributing to tissue injury and vascular permeability, more commonly reported in children^39^. Thus, while functional validation is awaiting, these findings combined with our cell-cell communication analysis reveal that imbalanced NK cell responses with reduced cell killing yet increased inflammation and migration may be implicated in SD progression.

Cytokines are considered key mediators of SD pathogenesis^40^. Among the 27 differentially secreted cytokines we identify between the same SDp and D patients analyzed via viscRNA-seq 2 were CCL2 (MCP-1), TNF, IL10 and IFNγ, previously implicated in increased plasma leakage and hepatic dysfunction in another pediatric cohort^41^. Interestingly, by integrating proteomic and transcriptomic data we identify candidate sources and interacting receptors of these cytokines. The increased serum MCP-1 level and upregulation of CCL2 and CCR2 transcripts we measure in myeloid cells from SDp may be associated with disease pathogenesis in humans, similarly to mice where CCR2-/- deletion reduces DENV-induced mortality and liver damage^42^. Increased serum TNF and upregulation of TNFα and TNFRSF1A-B transcripts in myeloid cells of SDp reveal activation of the TNFa pathway, previously shown to mediate tissue injury via caspase-3-mediated cell death^43^ and correlate with SD^44^.

High serum concentration of IL10—a suppressant of immune responses via interference with the NF-KB pathway, downstream IFN responses, and MHC-II expression^45^—has been implicated in SD^41, 46^ and shown to suppress DENV-specific responses upon restimulation of patient-derived T cells^47^. We measure higher levels of serum IL10 in SDp than dengue that is primarily driven by Tregs, and high IL10RA/B transcripts in myeloid and other cell types.

An intriguing hypothesis is that the activation of this pathway is involved in downregulating antigen presentation and IFN responses in APCs of SDp, thereby suppressing antiviral responses.

SDp show increased levels of IFNγ in serum (as previously reported^48^) largely derived from T and NK cells based on the viscRNA-seq 2, and increased IFN signaling genes (e.g. STAT1, IRF) in multiple cell types. Nevertheless, antigen presentation, effector responses, and ISGs—functions stimulated by IFNγ^49^—are suppressed rather than increased in APCs, consistent with impaired IFN responses. NK cells from SDp also downregulate IFN response genes, yet IFNγ may regulate the observed adaptive-like cytotoxic NK cell phenotype.

While non-neutralizing antibodies are key mediators of SD immunopathogenesis, we provide evidence that beyond ADE, functional and phenotypic alterations within specific cell types and subtypes affect the collective coordination and effectiveness of the immune system.

Such alterations may result from DENV-mediated immunomodulation, as observed by the downregulation of effector genes (IFN responses, MHC-II) in vRNA-harboring vs. bystander cells. Additionally, patient-specific attributes that interfere with antigen presentation, e.g. prior infections and genetic or epigenetic determinants, may predispose individuals to disease progression.

This study illustrates the utility of a systems immunology approach integrating viscRNA-Seq 2, flow cytometry, cytokine measurements and functional assays for understanding the immune response to viral infection. One limitation of this study is the focus on early infection. Future studies will assess the transcriptional alterations we detected in longitudinal samples, as we have recently done at the protein level^17^. Driven by the low number of PBMCs obtained from children, another limitation is the utilization of DWS samples for functional experiments.

Together, our systems immunology approach defined DENV target cells in the human blood providing evidence for active replication in B cells and discovered key correlates of disease progression in children, beyond ADE, with implications for prediction and prevention.

## Acknowledgements

This work was supported by an Investigator Initiated Award number W81XWH1910235 from the Department of Defense (DoD) office of the Congressionally Directed Medical Research Programs (CDMRP)/Peer Reviewed Medical Research Program (PRMRP), Catalyst and Transformational Awards from Dr. Ralph & Marian Falk Medical Research Trust, a National Institute of Allergy and Infectious Diseases (NIAID) grant U19 AI057229 supplement to S.E. and fuds from the Chan Zuckerberg Biohub to S.Q. and S.E.. S.E. is a Chan Zuckerberg Biohub investigator, who is also supported by a NIAID grant RO1AI158569, an Investigator Initiated Award number W81XWH2210283 and an expansion award number W81XWH2110456 from the DoD office of the CDMRP/PRMRP, and a Defense Threat Reduction Fundamental Research to Counter Weapons of Mass Destruction grant HDTRA11810039. L.G. was supported by EMBO Postdoctoral Fellowship ALTF 584-2021. Z.Y was supported by a Thrasher Research Fund early career award program grant and by a postdoctoral fellowship from the Maternal and Child Health Research Institute, Lucile Packard Foundation for Children’s Health. V.D. was supported by a Chan Zuckerberg Biohub Collaborative Postdoctoral Fellowship. M.L.R. was supported by the A.P. Giannini Foundation Postdoctoral Fellowship and the Harold Amos Medical Faculty Development Program. I.O. was supported by a Sue Merigan Student Scholar Fund in Infectious Diseases and Geographic Medicine. The funders had no role in study design, data collection and analysis, decision to publish, or preparation of the manuscript. We thank Dr. Catherine Blish for her guidance in the interpretation of NK cell data. Co-first authors L.G., Z.Y., Y.X. and V.D. contributed equally to this manuscript and each has the right to list themselves as first authors in their CVs.

## Author Contributions

Conceptualization: Y.Z, L.G, Y.X, V.D, S.E, F.Z; Validation: L.G, V.D, S.E; Methodology: Y.Z, L.G, Y.X, V.D, I.O, M.L.R, M.K.S, B.A.P, J.S, F.L, F.Z, S.E; Software: Y.Z, Y.X, L.G, V.D,

H.B.C, J.S, F.L, F.Z; Formal Analysis: Y.Z, Y.X, L.G, V.D, H.B.C, F.Z, S.E.; Investigation:

L.G, Y.Z, Y.X, V.D, S.E, F.Z; Resources: O.L.A.R, A.M.S, R.M.G, N.B, M.I.E.C, L.A.V.C,

E.M.R.G, F.R, M.L.R, J.G.M; Writing – Original Draft: L.G, S.E, Y.Z, V.D, Y.X, F.Z; Writing – Review and Editing: L.G, S.E, Y.X, V.D, F.Z; Supervision: S.E, F.Z; Funding Acquisition: S.E, F.Z, S.R.Q.

## Methods

### Colombia Cohort

#### Ethics Statement

All work with human subjects was approved by the Stanford University Administrative Panel on Human Subjects in Medical Research (protocols #35460 and #50513) and the ethics committees in biomedical research of the Fundación Valle del Lili (FVL, Cali, Colombia) and Centro de Atención y Diagnóstico de Enfermedades Infecciosas (CDI, Bucaramanga, Colombia). The parents or legal guardians of subjects provided written informed consent, and subjects between 2 to 17 years of age provided assent. Subjects were not involved in previous procedures and were all test-naïve. The demographics and clinical parameters of the subjects are summarized in **Table 1**.

#### Study Population and Sample Collection

The Colombia cohort consists of individuals presenting to the emergency room or clinics of FVL or CDI between March 2016 and Jan 2020. Enrollment criteria consisted of: i) age equal to or greater than 2 years; ii) presentation with an acute febrile illness of less than 7 day duration associated with one or more of the following symptoms or signs: headache, rash, arthralgia, myalgia, retroorbital pain, abdominal pain, positive tourniquet test, petechiae, and bleeding; and iiia) a positive dengue IgM antibody and/or NS1 antigen by the SD BIOLINE Dengue Duo combo device (Standard Diagnostic Inc., Korea) test or iiib) clinical presentation highly consistent with dengue and subsequent confirmation of diagnosis via serological testing and rRT-qPCR at Stanford. Patients were classified by infectious diseases specialists, both upon presentation and at recovery, as having dengue (D), dengue with warning signs (DWS), or severe dengue (SDp) according to 2009 WHO criteria^3^ (**Tables 1, 2**). Venous blood samples were collected upon enrollment on the first day of presentation. 10-40 ml of whole blood were collected in 1-4 tubes. Serum samples were obtained for additional assays. Sample transport, reception, and processing were strictly controlled using personal data assistants (PDAs) with barcode scanners.

For patients managed in an outpatient setting, follow-up was conducted daily via phone, during which patients were provided information about the clinical warning signs and asked about their appearance, until full recovery when final diagnosis was determined. Organ damage was defined according to standard clinical endpoints for DENV infection^50^. Demographics and clinical information were collected at the time of presentation (**Table 2**). The first day of fever (fever day 0) was defined by the patients or their relatives. Symptoms, warning signs, and laboratory parameters (including complete blood count, chemistry, and liver function test results) were documented by healthcare professionals.

For this study, PBMCs from 20 children infected with DENV (10 males and 10 females ranging, age: 4-17 years) and 4 healthy controls (2 males and 2 females, age: 5-12 years) were selected for viscRNA-seq and cytokine analysis (**Table 1**). Samples from 11 individuals were used for validation assays.

### Establishment of Dengue Diagnosis

#### Detection of DENV NS1 antigen and IgG/IgM

The SD BIOLINE Dengue Duo combo test (Standard Diagnostic Inc., Korea) was used to identify dengue patients for enrollment to the study.

#### rRT-PCR assays for detection of DENV and other microbial pathogens

To confirm the diagnosis of dengue and differentiate from infection with the co-circulating arboviruses, Zika virus and chikungunya virus, serum samples were screened with a qualitative, single-reaction, multiplex real-time reverse transcriptase PCR (rRT-PCR) that detects Zika, chikungunya, and DENV RNA. To identify the specific DENV serotype and quantify the viral load, samples positive for DENV in the screening assay were serotyped and quantitated using a separate DENV multiplex RT-qPCR^51^.

#### DENV serological assays

Anti-DENV IgG were tested using DENV Detect™ IgG ELISA kit (InBios, Seattle, WA) as per manufacturer instructions. Briefly, serum samples were diluted 1:100 in sample dilution buffer and 50uL was added per well. The top half (4 wells on each 8-well strip) of the ELISA plates were pre-coated with DENV derived recombinant antigen (DENRA) and the other half with normal cell antigen (NCA). Each sample went into two pairs of DENRA or NCA coated wells, one without and another containing 50 uL of 8 M Urea. Each plate contained a negative and a positive control provided in the kit. Plates were incubated in 37 °C for 1 hour and washed in a plate washer (Biotek 404 Select Microwasher). Next, we added 50 uL/well enzyme conjugate-HRP tagged goat anti-human IgG, incubated for 1 hour at 37 °C, and followed by wash. Next, 150 uL of EnWash was added, incubated for 5 min at room temperature and washed. Finally, 75 uL of TMB solution was added, incubated at RT in the dark for 10 min. The reaction was stopped by adding 50 uL stop solution. The plates were read at 450 nM by Spectra Max M2 (Molecular Devices) using SoftMax pro 7.0.3 software. Presence or absence of DENV IgG was interpreted from the ratio of readings from DENRA and NCA wells without urea. A ratio >=2.84 was considered positive, 1.65-2.84 ‘equivocal’ and <=1.65 as negative. The equivocal samples were repeated once. DENV IgG avidity was calculated by the ratio of reading in DENRA well without urea over the DENRA well with 8 M urea. High avidity (>0.6) was considered as secondary infection, moderate avidity (0.4-0.6), and low avidity (<0.6) as primary infection, in samples that showed a positive result by DENRA/NCA ratio.

#### PBMC isolation

PBMCs were isolated using SepMate tubes (Stemcell Technologies) according to the manufacturer’s instructions. Briefly, whole blood was diluted 1:1 with phosphate-buffered saline (PBS) and added to a SepMate tube, which contained 15 ml of Ficoll. Tubes were then centrifuged for 10 minutes at 1,200g, after which the PBMC layer was poured off into a fresh tube and washed with PBS. Tubes were then centrifuged at 250 x g for 10 minutes and resuspended in freezing media. Cryovials containing PBMCs were then placed in a CoolCell at -80 º C for 24 hours prior to being transferred to liquid nitrogen for storage.

#### Magnetic separation and rebalancing of PBMCs

Samples containing ∼1 ml of cryopreserved PBMCs were quickly thawed in a water bath at 37 ºC in medium containing 10% DMSO. Nine ml of warm medium were added and cells were spun at 300 g for 8 minutes. The supernatant was discarded, and 2 ml of medium were added. The samples were filtered with 35 μm cell strainers to obtain a pure single-cell suspension. The samples were mixed with magnetic microbeads to label cell debris, dead cells, and dying cells (Miltenyi Biotec) and the mix was set on a magnet to elute only living cells. The flow through was spun, the supernatant was removed, and the cell pellet was resuspended in 150 μl cell resining buffer with 0.5% BSA (AutoMACS). To enrich for rare cell populations, the samples were split into 2 aliquots: a 30 μl aliquot was kept intact, and a second aliquot was labeled with anti-CD3-Phycoerythrin (PE) antibody (BioLegend) and mixed with anti-PE magnetic microbeads to reduce the number of T cells. Cell numbers in both aliquots were measured, and the two aliquots were remixed. The samples were then diluted to 1000∼1400 cells/μl and processed via viscRNA-seq 2.0.

#### Virus-inclusive single cell RNA-sequencing (viscRNA-seq 2)

The master mix containing 10,000-14,000 cells per sample was prepared following the standard 10X Genomics 5′ RNA-Seq protocol. A DENV capturing oligo was added to the master mix at a final concentration of 30 nM for capturing the specific DENV serotype identified via RT-qPCR in patient’s sera (DENV1: 5′-AAGCAGTGGTATCAACGCAGAGTACTTTCCCCAGCTTTTCCATGA-3′); DENV3: 5′-AAGCAGTGGTATCAACGCAGAGTACTTTCCCCACGTTTTCCATGA-3′).

Samples were loaded into the droplet generator for partitioning single cells into an inverted emulsion. The Next GEM Single Cell 5’ reagent kit (10x Genomics) was used for reverse transcription, cDNA amplification and construction of gene expression libraries. Library quality was assessed using a 2100 Bioanalyzer High Sensitivity DNA kit (Agilent).

Sequencing was performed in a paired-end mode with S1 and S4 flow cells (2 × 150 cycles) on a NovaSeq 6000 sequencer (Illumina).

#### scRNA-seq data analysis

Python 3 and Jupyter notebooks were used for the analysis and are available at https://github.com/saberyzy/DENV. The following open source software was used for this study: numpy^52^, pandas, matplotlib, seaborn, anndataks 0.1.3, pysam and scanpy.

#### Pre-processing of 10x Genomics Chromium scRNA-seq data

CellRanger v3.1.0 (10x Genomics) was used to process the raw read pairs. To generate a gene expression matrix for each sample, we constructed a combined reference of (i) human genome (GRCh38) and (ii) DENV genome as an additional chromosome (GenBank GQ868569 for DENV 1 and MH544647 for DENV 3). We then aligned the reads to this reference and measured the number of UMIs for each host gene and vRNA in each cell. High quality cells were scored based on the following criteria: fewer than 15% mitochondrial reads; over 400 expressed genes; fewer than 30,000 WHAT??; and over 1,000 detected UMIs. We excluded genes that were expressed in less than three cells. The resulting dataset was normalized by the total UMI count, multiplied by 1,000,000 (cpm) and log2 transformed after addition of a pseudo count of 1.

#### Dimensional reduction, clustering and cell type annotation

After normalization, principal component analysis (PCA) was performed on the 2,000 most variable features identified by dispersion-based methods. The normalized dispersion was obtained by scaling with the mean and standard deviation of the dispersions for genes falling into a given bin for mean expression of genes. The two-dimensional embedding was computed via UMAP based on the first 15 dimensions of the PCA reduction. The cells were clustered using the Leiden algorithm based on the first 15 PCA dimensions with modularity metric and a resolution parameter of 0.3. Clusters were annotated based on marker genes from public sources to identify immune cell types and were confirmed using Azimuth. The data from each annotated immune cell type were further embedded and clustered to annotate cell subtypes.

#### Differential gene expression analysis and gene enrichment analysis

DEGs identified by a 2-sample Kolmogorov-Smirnov test using anndataks 0.1.3. Genes with >1 log2 fold change between the two groups and a p-value <=0.05 (after FDR correction for multiple hypotheses) were selected for downstream analyses. Metascape and GSEAPY were used for pathway analysis of groups of up- or downregulated genes.

#### Detection of vRNA harboring cells (VHCs)

We counted the number of molecules aligning against the DENV chromosome in our composite reference. All cells with >= 1 detected viral molecules were considered VHCs, else they were considered bystanders. Since the number of viral molecules detected in a single cell was never above 24 and typically 1 or 2, the likelihood of vRNA spillover from separate droplets is low. Several analyses were carried out using slightly different thresholds for VHC versus bystander and yielded similar results for the fraction of positive/negative strand reads.

#### Machine learning prediction of SD progression based on cell subtype abundance

To predict patient progression to SD based on cell subtype abundance, we computed the fraction of proliferating plasmablasts, signaling NK cells, and non-classical monocytes in each patient as a three-dimensional vector and trained a support vector machine (SVM) regression model with a third-degree polynomial kernel using the class NuSVC in scikit-learn. We chose SVMs partly because they have a straightforward geometrical interpretation as one can directly plot the hypersurface with the nullcline of the decision function (black dashed curve in **Fig. 3c** and gray surface in **Fig. 3d**). We assessed the performance of the model via leave-one-out validation on all patients.

#### Cell-cell communication analysis

We extracted the list of known potential interaction partners from OmniPath and complemented it with a series of interactions mined from recent immunology literature (**Table 6**). Given the shallow sequencing of droplet single cell platforms, interactions were counted if both the ligand and the receptor were expressed in more than 2 % of the cells within the respective cell type. Variations in this threshold did not affect our results. Interacting DEGs (iDEGs) were defined as ligands or receptors in our list of interactions which were clear DEGs (i.e. |log2-fold change| > 1 and an up- or - downregulation in at least 40 out of 54 pairwise patient comparisons) (**Fig 6.b, top**).

Differential expressed interactions (DEIs) between D and SDp were classified as upregulated and downregulated if both the receptor and ligand were overexpressed or under-expressed, respectively, or as inversely regulated if one of the partner genes was overexpressed and the other under-expressed (**Fig 6.b, bottom**). A nonparametric label randomization test was utilized to define the significance of differential interactions. The p-value was calculated as the fraction of randomizations for which the distance from the origin was larger (i.e., differential expressions were more extreme) than in the actual data (**Extended Data Fig 6.g**). Violin plots were designed to show the expression of the interacting gene on the corresponding cell type in each patient to verify the robustness of DEIs passed the randomization test (**Extended Data Fig 6.h**).

#### Spectral flow cytometry

Cryopreserved PBMCs were thawed at 37 °C, washed, and incubated with the Zombie Aqua™ live/dead fixable dye (Biolegend) at 1:1000 and with 5 μl of FcR blocking reagent (Miltenyi) for 15 min at room temperature. Cells were labeled for 20 min at 4 °C with the following specific antibodies: BUV496 anti-CD19 (SJ25C1;BD), BUV563 anti-CD56 (MY31;BD), BUV615 anti-HLA-DR (G46-6;BD), BUV661 anti-CD20 (2H7;BD), BUV737 anti-CD16 (3G8;BD), BUV805 anti-CD3 (SK7;BD), BV421 anti-CD1c (L161;Biolegend), BV605 anti CD11c (3.9;Biolegend), BV711 anti CD163 (GHI/61;Biolegend), PeCy5 anti-CD123 (6H6;Biolegend), PeCy7 anti-CD141 (M80’;Biolegend), APC anti CD14 (M5E2;Biolegend), AF700 anti CD303 (201A;Biolegend), APC-Cy7 anti CD15 (W6D3;Biolegend) and PeCy5.5 anti-Flavivirus (4G2;Novus Biologicals). Cells were fixed with 1% PFA for 24 h, and data were acquired on an Aurora spectral analyzer (Cytek) and analyzed with the FlowJo software (Tree Star).

#### Co-culture experiments assessing infectious DENV production

Cryopreserved PBMCs were thawed at 37 °C and washed with cell culture media (RPMI (Gibco) supplemented with 10% FBS + 20 U/ml sodium heparin (Sigma) and 0.025 U/ml Benzonase (Sigma). Different cellular fractions were sorted via fluorescence activated cell sorting (FACS, BD ARIA) after staining with FITC anti-CD45 (HI30;Biolegend), BV605 anti-CD3 (OKT3;Biolegend), PE-Cy5.5 anti-CD19 (HIB19;BD), PE anti-CD20 (2H7;Biolegend), and APC-Cy7 anti-HLA-DR (L243;Biolegend) antibodies. 2×10^4^ sorted cells were added to 4×10^4^ human hepatoma cells (Huh7) and incubated for 96 hours at 37 °C. Co-cultured cells were then lysed and RNA extracted with RNeasy kit following manufacturer′s instructions. DENV RNA copies number were determined via RT-qPCR.

#### Uptake assay

Cryopreserved PBMCs were thawed at 37 °C, washed with cold cell culture media (300 g, 4 °C) and pre-incubated for 30 min either at 4 °C or at 37 °C. 5×10^5^ cells were then incubated with 1 mg/ml of pHrodo-labeled *S*.*aureus* bioparticles (ThermoFisher) for 1 hour at 4 °C to determine non-specific binding on the cell surface (binding control), or at 37 °C for active uptake. Cells were harvested, labeled with fluorescently-labeled antibodies as described above, and pHrodo fluorescence was assessed by spectral flow cytometry.

#### Cytokine measurement

Cytokine concentration from DENV infected patients’ serum samples were determined via Luminex Assay Human 80 (R&D System) following manufacturer′s instructions (**Table 5**).

#### Cytokine-receptor communication

The serum concentrations of cytokines of SDp and D patients were compared by Man-Whitney U test. Differentially secreted cytokines were defined as those with a p value < 0.05. The RNA expression of cytokine producing genes and corresponding receptors was then computed in distinct cell subtypes (**Figure 6H, Figure S6I**).

## List of Tables

**Table 1**: Patients ID and clinical data

**Table 2**: Number of high-quality cells for each patient

**Table 3**: Differentially expressed genes between SDp and D patients

**Table 4**: Candidate cell-cell interactions in SDp vs. D patients

**Table 5**: Cytokine concentration in patients serum

**Table 6**: Additional cell-cell interactions – source table

