## Supplementary Figures for "Global and cell type-specific immunological hallmarks of severe dengue progression" (from another version of this preprint: version 1, 2022-12-14)

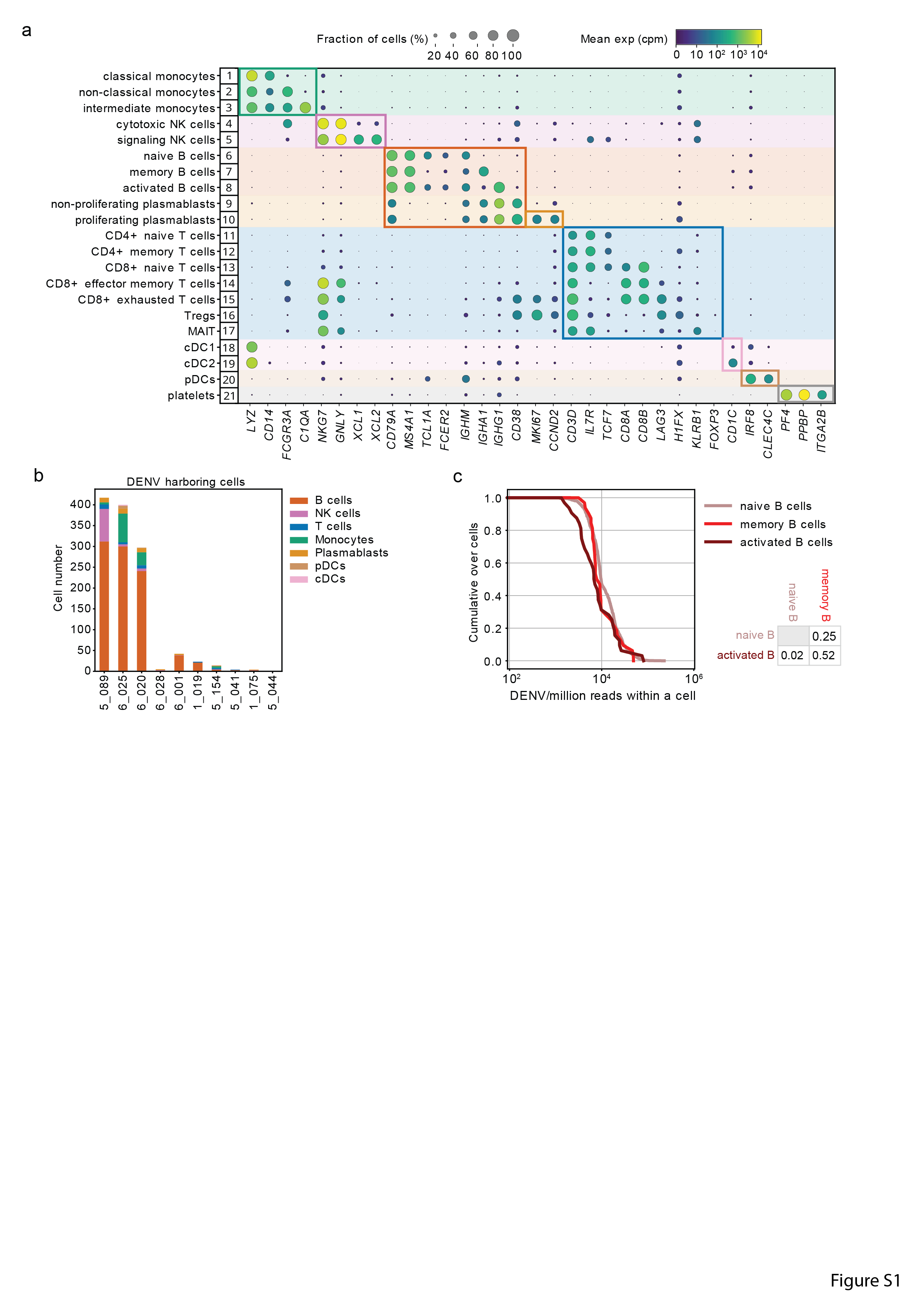

**Fig.1: PBMCs annotation and identification of VHCs via viscRNA-Seq 2. a**, Dot plot depicting examples of marker genes used to annotate the indicated 21 immune cell subpopulations in the visc-RNA seq dataset (Fig.1a). Dot size indicates the fraction of cells expressing the marker gene; color indicates expression level of the marker gene in cpm; and identification numbers of distinct cell populations refer to the UMAP in Fig.1a. **b**, Stack bar plots showing the absolute numbers and distribution of VHCs across cell types in each of the 10 patients with detectable vRNA. **c**, Cumulative distributions of DENV reads/million reads per cell in B cell subtypes. values indicate p-values from ks test after Bonferroni correction between distinct B cell subtypes.

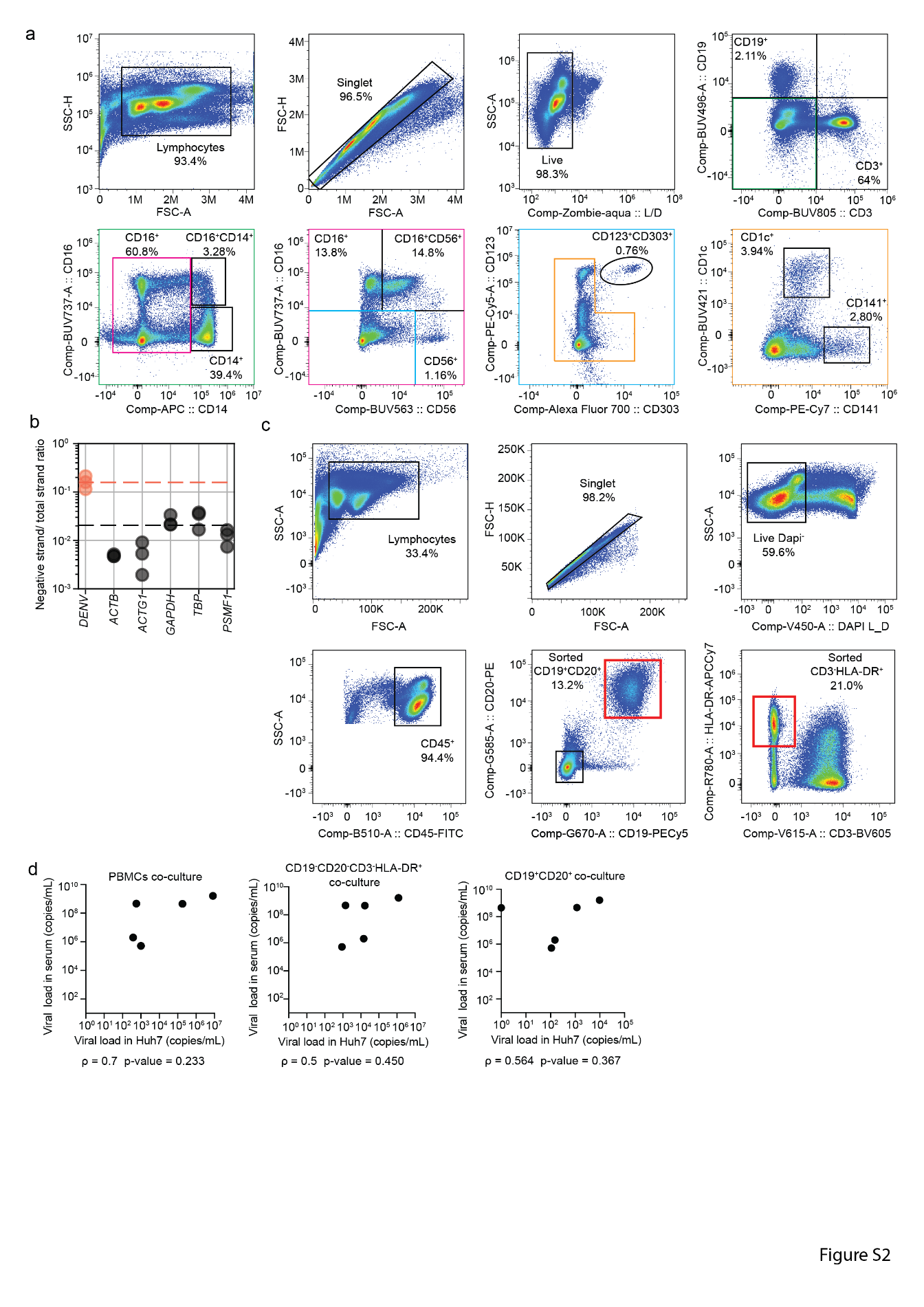

**Fig.2: B cells from DENV-infected patients harbor replicating virus. a**, Flow cytometry gating strategy used to define B cells (CD19^+^), T cells (CD3^+^), classical monocytes (CD14^+^CD16^-^), non-classical monocytes (CD14^-^CD16^+^), intermediate monocytes (CD14^+^CD16^+^), CD56^bright^ NK cells, CD56^dim^CD16^+^ NK cells, pDCs (CD123^+^CD303^+^), cDC1s (CD141^+^) and cDC2s (CD1c^+^). **b**, Dot plot showing the average ratio of negative strand over total strand of housekeeping genes (gray) and DENV RNA (orange) reads within the 20 single cells with the largest DENV read counts in 3 DWS patients with highest vRNA reads shown in **Fig.1e**. **c**, FACS gating strategy used to sort populations of B cells (CD19^+^, CD20^+^) and CD19^-^CD20^-^ CD3^-^HLA-DR^+^ cells for co-culture experiments. **d**, Scatter plots showing correlations between viral load in serum and viral load in Huh7 cell lysates (DENV copies /mL) following 5-day co-culture either with PBMCs or with CD19^-^CD20^-^ CD3^-^HLA-DR^+^ and B cell (CD19^+^, CD20^+^) fractions from the same patients. Spearman correlation coefficients and p-values are shown below each panel.

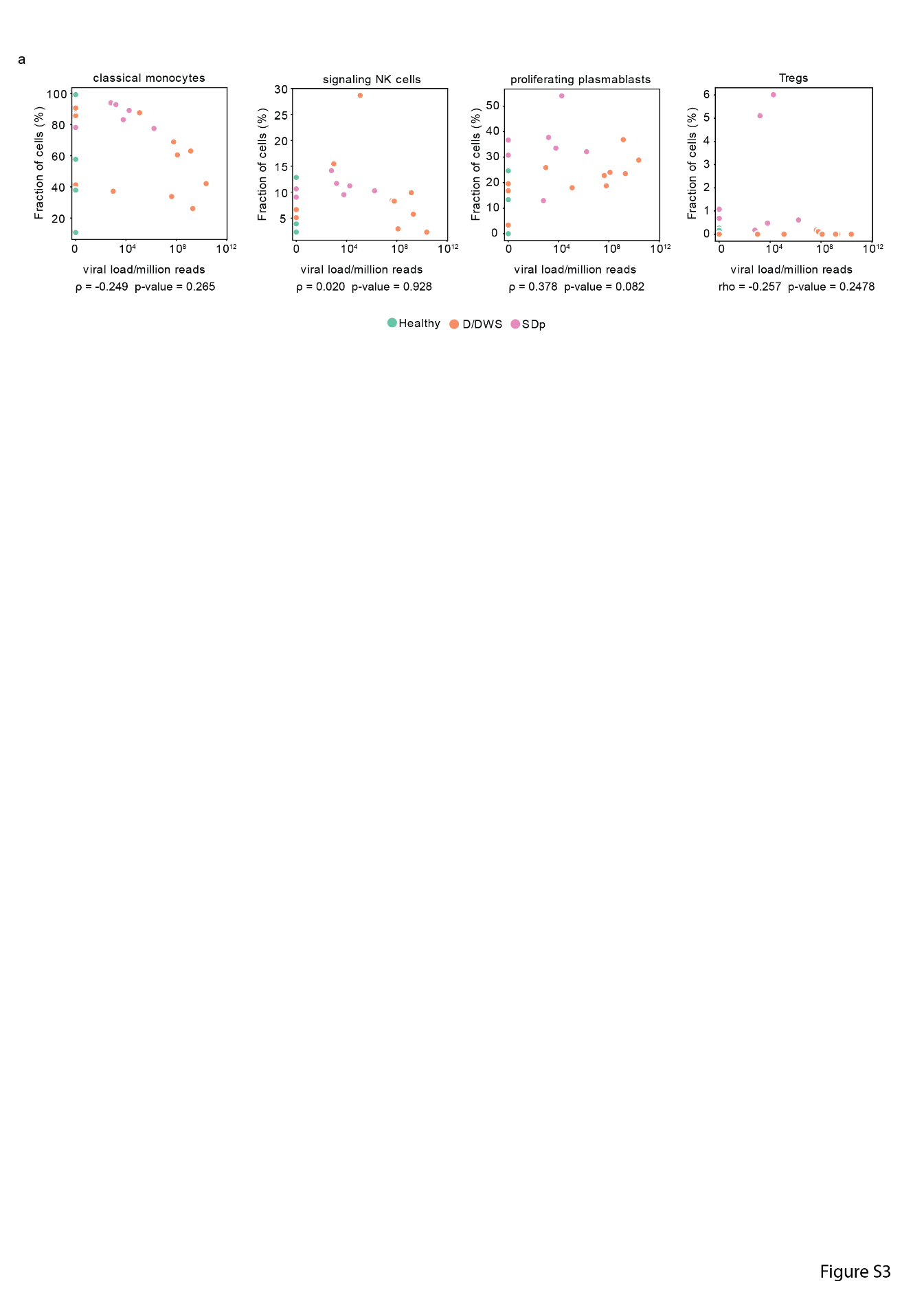

**Fig.3: Differences in cell subtype abundance do not correlate with serum viral load. a**, Scatter plots showing correlations between fractions of classical monocytes, signaling NK cells, proliferating plasmablasts, and Tregs and serum viral load measured by RT-qPCR. Each dot represents a single patient colored by disease severity (H= green; D/DWS= orange; SDp= pink). Spearman correlation coefficients and p-values are shown below each panel.

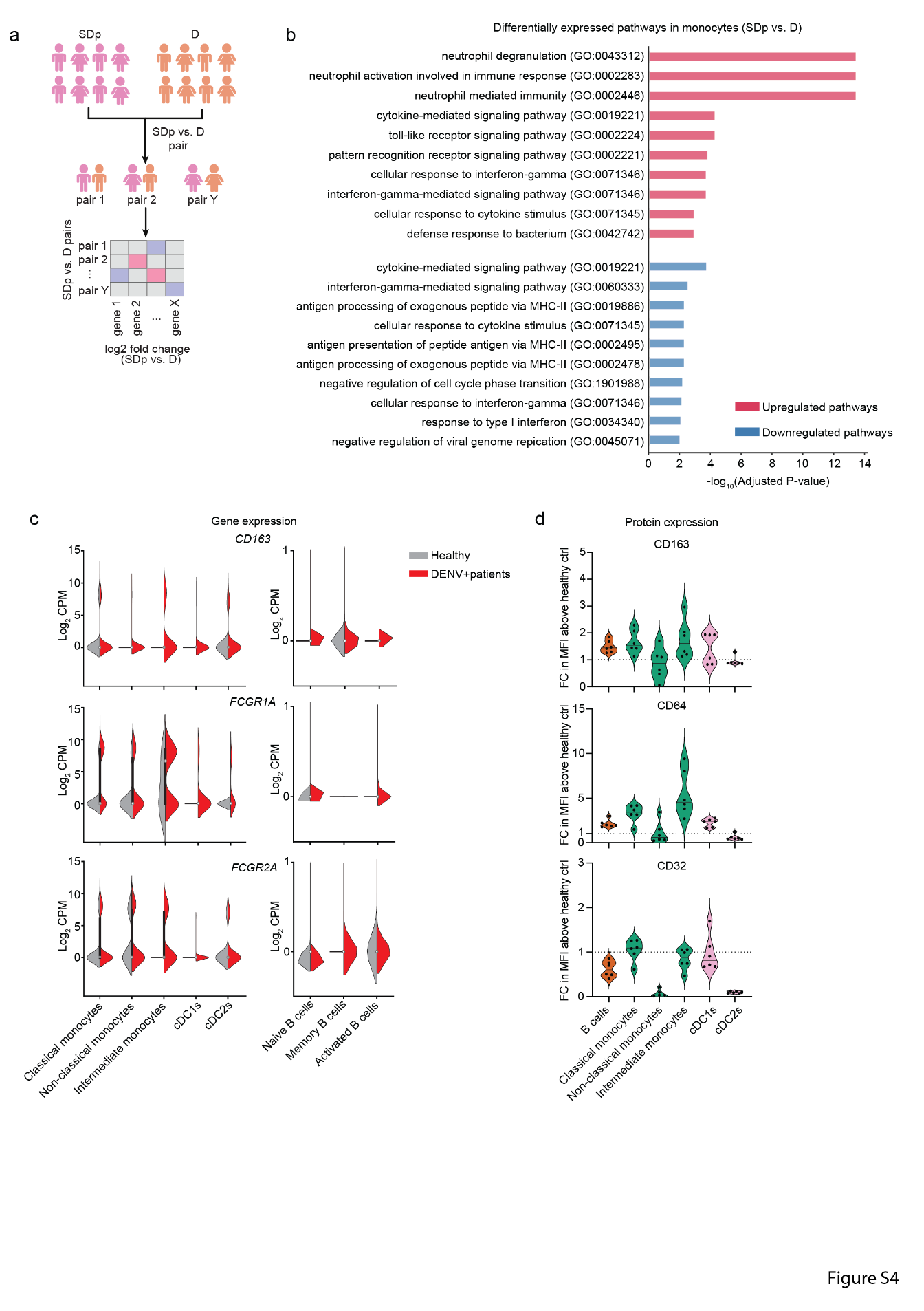

**Fig.4: APCs from SDp show signatures of increased activation but decreased antigen presentation. a**, Schematic illustrating the pairwise comparison strategy used to identify DEGs between SDp and D patients. Pairs were generated considering only patients that showed n≥5 cells in a cell type or n≥3 in a cell subtype. For each gene, we calculated the geometric mean expression in various cell subtypes for each patient in the pair and the log2 fold change in expression between the two patients in the pair. A median log2 fold change was then obtained for each gene by analyzing all pair combinations. DEGs were defined as those with a median log2 fold change greater than 1 or smaller than -1. **b**, Pathway analysis showing top 10 upregulated and downregulated gene ontology (GO) terms between SDp and D in monocytes. **c**, Violin plots showing expression (Log2 CPM) of CD163, FCGR1A and FCGR2A in monocytes, cDCs and B cell subtypes in healthy (gray) and DENV+ patients including SD and D (red). **d**, Violin plots showing expression of CD163, CD64 and CD32 proteins on B cells (CD19^+^), classical monocytes (CD14^+^CD16^-^), non-classical monocytes (CD14^-^CD16^+^), intermediate monocytes (CD14^+^CD16^+^), cDC1s (CD141^+^) and cDC2s (CD1c^+^) measured via spectral flow cytometry in DWS patient-derived PBMCs. Data are shown as fold change (FC) of mean fluorescence intensity (MFI) above healthy control (n=6, N=2). Dotted line represents mean value of healthy controls normalized at value 1, horizontal line in each violin plot represents median value.

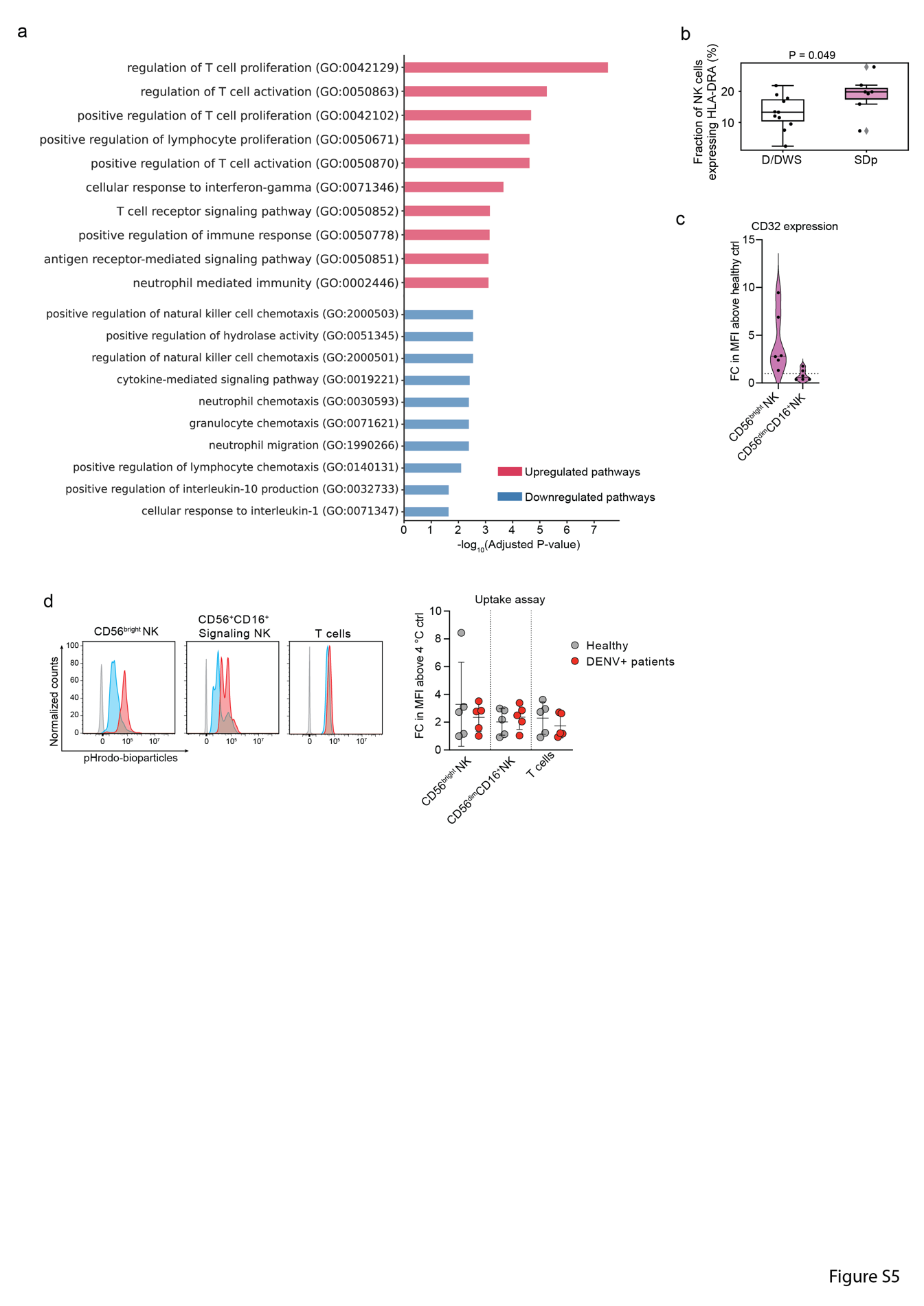

**Fig.5: NK cells in SDp show activation and adaptive-like signatures. a**, Pathway analysis showing top 10 upregulated and downregulated gene ontology (GO) terms between SDp and D in NK cells. **b**, Fraction of NK cells expressing HLA-DRA gene out of the total NK cell population in D/DWS and SDp. **c**, Violin plots showing CD32 protein expression in CD56^bright^ NK cells and CD56^dim^CD16^+^ NK cells measured via spectral flow cytometry in patient-derived PBMCs. Data are shown as fold change (FC) of mean fluorescence intensity (MFI) above healthy control (n=6, N=2). Dotted line represents mean value of healthy controls normalized to 1; horizontal line in each violin plot represents median value. **d**, Histograms showing the percentage of pHrodo Bioparticles positive cells in distinct cell subtypes derived from healthy (gray) and DENV-infected patients measured at 4 °C (cyan) or 37 °C (red). Dot plot showing bioparticle uptake quantification as FC of mean MFI above 4 °C control. Each dot represents an individual color-coded by disease status: healthy (gray) and DENV-infected (red) (n=5, N>2).

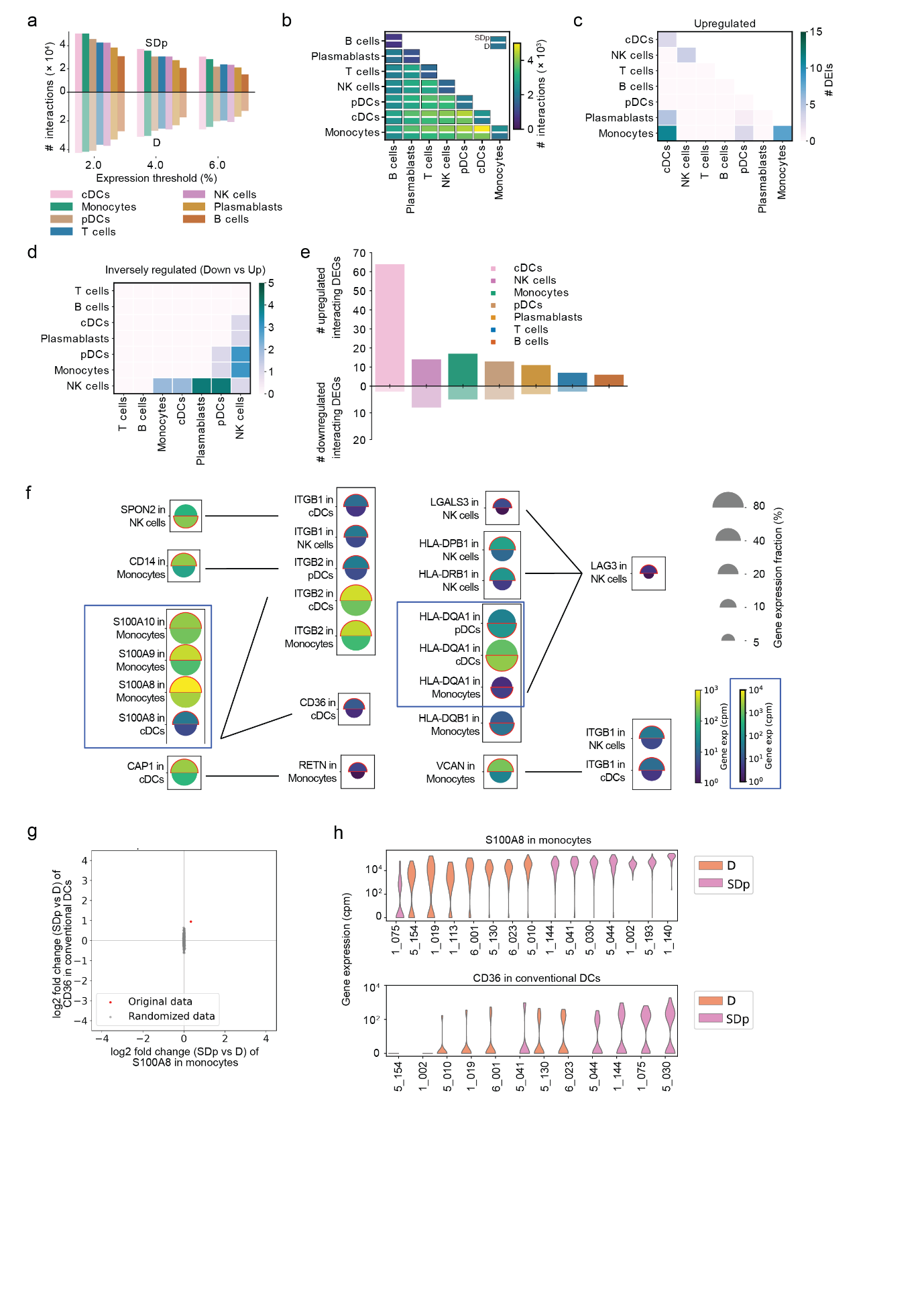

**Fig.6: Cell-cell communications are altered in SDp. a**, Bar plot showing the number of candidate interactions in each cell type in SDp (top) and D (bottom) based on different expression thresholds (2%, 4%, and 6%). **b**, Heatmap showing the number of candidate interactions between the indicated cell types in SDp (top rectangle) and D (bottom rectangle). **c, d,** Heatmaps showing the number of upregulated and inversely regulated DEIs shown in fig. 6c and 6d, respectively, across cell types. **e**, Bar plot showing the number of up- and down-regulated interacting DEGs across cell types in SDp vs. D. **f**, Split dot plots showing gene expression in representative DEIs. Top half of each spilt dot plot depicts the expression level in SDp; bottom half depicts the expression level in D; spilt dot plot size depicts the percentage of gene expressing cells in the cell types indicated; and color depicts gene expression level in counts per million (cpm). Red outline indicates the condition with the higher gene expression. **g,** Example of label randomization for the S100A8-CD36 interaction. The red dot indicates the log2 fold change of the expression of either gene in the corresponding cell type in SDp vs D. The 1000 gray dots indicate the log2 fold changes after randomly picking SDp and D cells following cell shuffling. The p-value was calculated as the fraction of randomizations for which the distance from the origin was larger than in the real data. **h**, Violin plots depicting ligand-receptor expression (S100A8-CD36 interaction) in each patient.
